# Genomic and Transcriptomic Insights into the Evolution and Parasitic Strategy of the Woody-Plant Nematode *Pratylenchus vulnus*

**DOI:** 10.1101/2025.09.24.678327

**Authors:** Dadong Dai, Yali Zhang, Romnick Latina, Xuyun Yang, Valerie M. Williamson, Simon C. Groen, Syed Shamsullah, Charles A. Leslie, Bardo Castro, Shahid Siddique

**Author notes:** Corresponding author: Professor Shahid Siddique. These authors contributed equally to this manuscript (Dadong Dai, Yali Zhang).

## Abstract

The root-lesion nematode *Pratylenchus vulnus* parasitizes a wide range of hosts including woody perennials such as walnut (*Juglans regia*) and grapevine (*Vitis vinifera*), significantly damaging roots and reducing yields. Here, we present a high-quality, chromosome-level genome assembly of *P. vulnus* (61.7 Mb across six chromosomes). Comparative genomic analysis revealed high collinearity in protein-coding genes between *P. vulnus* and the root-knot nematode *Meloidogyne graminicola*, indicating a closer evolutionary relationship with this sedentary endoparasite. Large chromosomal regions in *P. vulnus* lack synteny with other nematode genomes, have comparatively low GC content (<30%), and are enriched in genes with unique or lineage-specific functions. Transcriptome analysis highlighted dynamic, stage-specific expressions of genes involved in parasitism, development, and metabolism. Additionally, we identified an extensive repertoire of putative effector genes and characterized lineage-specific expansions of cell wall-degrading enzyme families. Overall, these findings provide insight into the genome organization, chromosome evolution, and parasitism-related gene repertoire in a woody-plant parasitizing nematode.

## Introduction

Plant-parasitic nematodes (PPNs) are among the most destructive plant pests, causing billions of dollars in crop losses globally each year (Jones et al., 2013). Among PPNs, root-lesion nematodes (RLNs) of the genus *Pratylenchus* are particularly damaging due to their broad host range, affecting both herbaceous and woody plant species (Fosu-Nyarko and Jones, 2016, Jones et al., 2013). The infection cycle of RLNs begins when second-stage juveniles (J2s) hatch from eggs deposited in the soil or in root tissues. Unlike sedentary endoparasitic nematodes, all life stages of RLNs - including third- and fourth-stage juveniles, as well as adult males and females - remain motile and can enter and leave host root tissues guided by root exudates and other environmental cues (Friebe et al., 1998, Fosu-Nyarko and Jones, 2016). Once inside, they move inter- and intracellularly through the root cortex, feeding on live cells with a stylet and causing characteristic necrotic lesions. The entire life cycle can be completed within a few weeks under favorable environmental conditions, allowing for multiple generations per growing season and contributing to their persistence and damage, especially in perennial cropping systems (Chihani-Hammas et al., 2018, Karaca et al., 2020, Fosu-Nyarko and Jones, 2016).

Among RLNs, *Pratylenchus vulnus* poses a significant threat to perennial crops such as walnut (*Juglans regia*), grapevine (*Vitis vinifera*), and various stone fruits (Karaca et al., 2020, Chihani-Hammas et al., 2018). In California, where walnuts are a major agricultural commodity, *P. vulnus* poses an existential threat to the industry (Pollack, 1998, McKenry, 1989). Although comprehensive statewide surveys are lacking, yield losses attributed to RLNs in walnut orchards are estimated to range from 10% to 20%, potentially translating to $140 million in annual losses (Personal Communication, California Walnut Board and Commission). In severe cases, yield reductions can reach 50% or more. Traditionally, management in walnut orchards has relied on soil fumigants such as 1,3-Dichloropropene. However, growing regulatory restrictions and environmental concerns have made these chemical control methods less viable, highlighting the urgent need for alternative, sustainable strategies.

Despite the agricultural importance of *P. vulnus*, genomic resources for this species—and RLNs more generally—remain extremely limited (Kikuchi et al., 2017). By comparison, substantial progress has been made in the genomic analysis of sedentary endoparasites such as root-knot nematodes (RKNs, *Meloidogyne* spp.) (Dai et al., 2023) and cyst nematodes (CNs, *Heterodera* and *Globodera* spp.) (Siddique et al., 2022), which primarily parasitize annual herbaceous crops. Interestingly, molecular phylogenetic analyses have shown that although both RKNs and CNs are sedentary PPNs, RKNs are more closely related to RLNs, whereas their evolutionary relationship with CNs is more distant (Qing et al., 2024). Genomic studies of RKNs have revealed complex ploidy differences, evidence of horizontal gene transfer (HGT), and effector diversification underpinning their parasitic strategies (Dai et al., 2023). However, the lack of comparable data for RLNs restricts our understanding of parasitism and the evolution of host adaptation in this group (Kikuchi et al., 2017).

Here, we present a chromosome-level genome assembly of *Pratylenchus vulnus*, providing a foundational genomic resource for RLNs. We integrated genome-wide analyses of gene content, synteny with other nematode species, and chromosome structure with comparative transcriptomic profiling to investigate host-induced gene expression dynamics and effector deployment.

## Results

### Chromosome-level assembly and annotation of the *P. vulnus* genome

To better understand the genomic features associated with the ability of migratory nematodes to parasitize woody plants with lignified xylem, we sequenced and assembled the genome of *P. vulnus* strain *Yolo1*. First, we generated 15 Gb of short-read Illumina sequencing data and analyzed it using Smudgeplot, confirming that *P. vulnus* is diploid (**Fig. S1A**). Using GenomeScope2, we estimated a genome size of approximately 57 Mb (**Fig. S1B**). For genome assembly, we utilized 20 Gb of long-read Nanopore data and 8 Gb of PacBio HiFi data, yielding a 61.7-Mb draft genome assembled into 63 contigs (**Table 1**). Subsequently, three rounds of polishing with Illumina short reads improved sequence accuracy. To assess potential contamination from bacterial or other non-target DNA sequences in our assembly, we performed BlobPlot analysis (Challis et al., 2020) of the assembled contigs. The results showed that 89.5% of the total length was classified as coming from Nematoda, 6.9% had no hits found (no-hit), and only three contigs, accounting for 3.6% of the total sequence length, were initially identified having non-nematode origins (**Fig. S2**). Upon closer manual inspection of the BLAST alignment details for contigs identified as non-nematode in origin, we found that only short, discontinuous regions of a few hundred base pairs aligned to potential contaminant taxa. Therefore, we consider these contigs to be more appropriately categorized as no-hit.

**Table 1:** Statistical summary for the *Pratylenchus vulnus* genome assembly and annotation.

| Assembly Feature |  | Annotation Feature |  |
| --- | --- | --- | --- |
| Contig-level assembly | Value | Genes | Number |
| Initial genome size (NECAT) | 61.72 Mb | protein-coding genes | 13,311 |
| Polished genome size | 61.67 Mb | transcripts (isoforms) | 18,704 |
| Number of contigs | 63 | transcripts with UTRs | 24,250 |
| Contig N50 | 6.6 Mb | Transcripts with Pfam domains | 12,444 |
|  |  | Transcripts with GO terms | 9,016 |
| **Chromosome-level assembly (Hi-C)** |  |  |  |
| Final genome size | 61.71 Mb | **Chromosome** | size |
| Number of scaffolds | 13 | Chromosome 1 | 14,202,844 bp |
| Number of chromosomes | 6 | Chromosome 2 | 12,102,447 bp |
| Scaffold N50 | 10.1 Mb | Chromosome 3 | 10,106,057 bp |
| Percentage of sequences anchored to chromosomes | 99.88% | Chromosome 4 | 8,859,133 bp |
|  |  | Chromosome 5 | 8,682,852 bp |
|  |  | Chromosome 6 | 7,683,060 bp |

To achieve a chromosome-level assembly, we performed chromatin conformation capture (Hi-C) sequencing, generating 78 Gb of data. Using chromatin interaction information, we anchored 88.89% of the contigs to six pseudochromosomes (**Fig. 1A**), consistent with previous karyotype studies on this species (Roman and Triantaphyllou, 1969). To assess the completeness of our assembled genome, we searched for tandem repeat sequences and identified from 67 to over 4000 copies of the typical telomeric repeat sequence TTAGGC at the ends of each chromosome (**Fig. 1B**). G-quadruplex motifs were unevenly distributed across the chromosomes (**Fig. 1B**), showing enrichment in some regions and scarcity in others, suggesting potential structural or regulatory roles in specific chromosomal domains (Du et al., 2009).

**Fig. 1:**
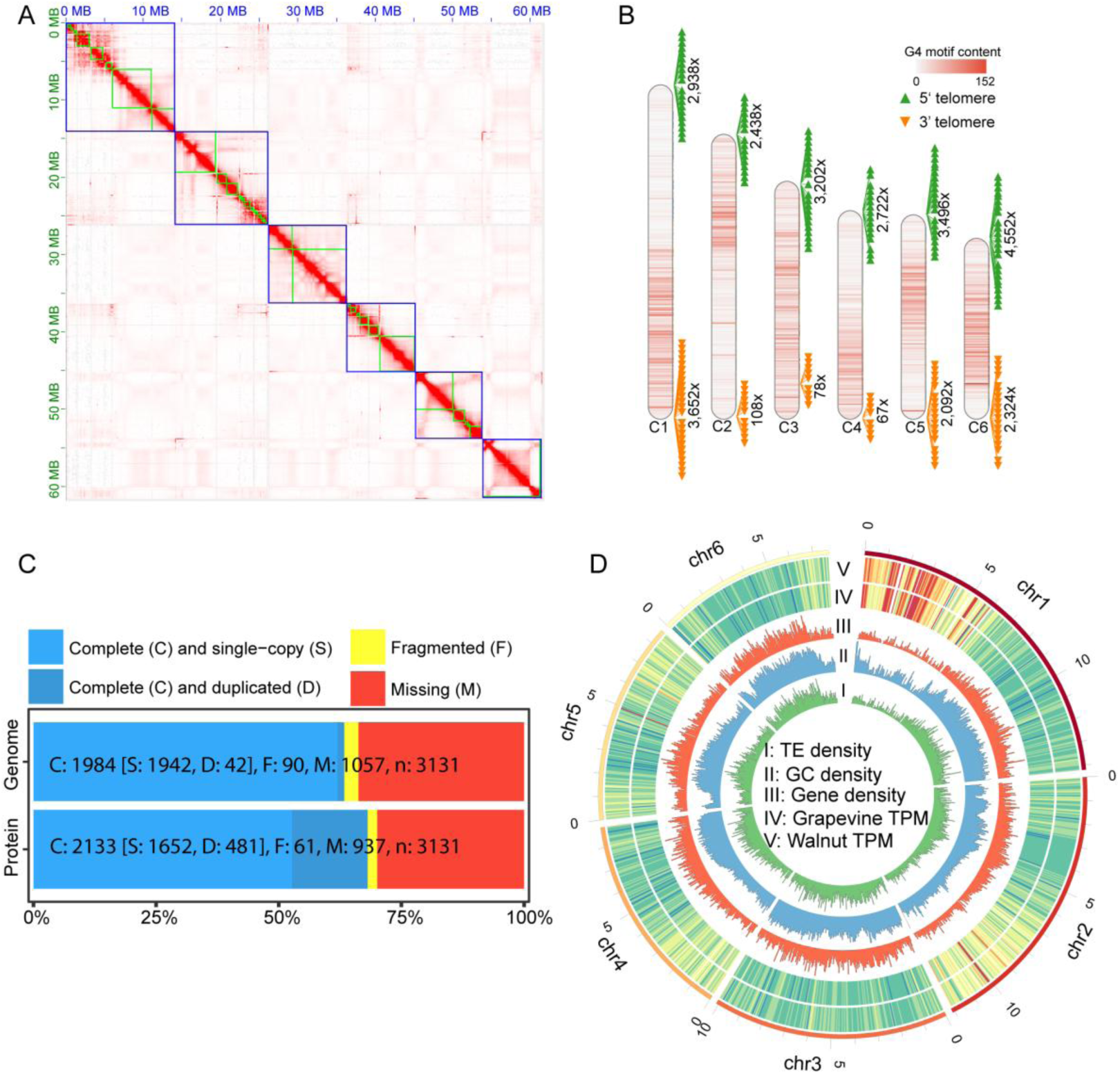
Overview of the *Pratylenchus vulnus* genome. **A.** Hi-C contact heatmap showing chromosome-scale assembly of the *P. vulnus* genome. The green boxes represent contigs, and the blue boxes represent pseudochromosomes. **B.** Chromosomal distribution of G4 DNA motifs and telomeric repeats (TTAGGC). Orange and yellow arrowheads mark the 5′ and 3′ telomeres, respectively. The heatmap shading indicates G4 motif content. Numerical values (×) indicate the number of occurrences of the telomeric repeat sequence at the corresponding position. **C.** BUSCO analysis showing genome completeness based on gene (top) and protein (bottom) datasets. Categories: complete and single-copy (light blue), complete and duplicated (dark blue), fragmented (yellow), and missing (red). **D.** Circos plot displaying genomic features across the six *P. vulnus* chromosomes, displayed in five concentric circles: I, transposable element (TE) density; II, GC content; III, gene density; IV, gene expression in transcripts-per-million (TPM) in *Vitis vinifera* (grapevine); V, gene expression (TPM) in *Juglans regia* (walnut). n circles IV and V, blue indicates high expression and red indicates low expression.

To support our gene predictions and functional annotation, we collected mixed-stage nematodes parasitizing the roots of walnut trees or grapevine and extracted total RNA for library construction and sequencing. We identified 13,281 protein-coding genes and 18,704 transcripts, including 5,423 alternative isoforms. Functional annotation using the Gene Ontology (GO), Kyoto Encyclopedia of Genes and Genomes (KEGG), and Pfam databases assigned putative functions to 77.2% (14,432) of the transcripts. Additionally, approximately 9.6% of the genome consisted of repetitive sequences, comprising 21,502 transposable elements (TEs; **Supplementary Table S1**). Annotating non-coding RNA-transcribing genomic loci revealed 371 tRNAs, 88 rRNAs, and 66 snRNAs. We also identified 712 transcription factor-encoding genes from 92 families, 1,629 genes encoding secreted proteins (candidate effectors), and 326 horizontally transferred genes.

We assessed genome completeness using Benchmarking Universal Single-Copy Orthologs (BUSCO) and achieved completeness scores of 63.3% for genomic sequences and 68.2% for protein sequences (**Fig. 1C**), comparable to other published PPN genomes(Dai et al., 2023). Finally, we used Circos plots (Krzywinski et al., 2009) to visualize the genomic landscape, showing gene density, gene expression levels, GC content, and TE density (**Fig. 1D**). Notably, the left arm of chromosome 1, the right arm of chromosome 2, and specific regions on chromosomes 4 and 5 exhibited lower gene density, lower TE and GC contents, and lower gene expression levels relative to the genome-wide averages (**Fig. 1D).**

### Developmental, host-related, and cold-induced gene expression signatures in *P. vulnus*

To explore transcriptomic variation across the nematode life cycle, we manually collected nematodes at six distinct developmental stages—eggs, second-stage juveniles (J2), third-stage juveniles (J3), fourth-stage juveniles (J4), males, and females—and performed deep RNA sequencing (RNA-seq) of each stage (**Fig. 2A**). In addition, to compare gene expression of nematodes from different hosts, we analyzed mixed-stages transcriptomes from walnut and grape. Furthermore, given that *P. vulnus* is known to survive extended periods of low temperatures typical of winter soil conditions commonly occur in Mediterranean-type climates like California’s Central Valley, we sought to investigate its early molecular response to cold exposure. To do this, we exposed a mixed-stage population to 4 °C for 6 hours to capture initial transcriptional changes triggered by acute cold treatment (**Fig. 2A**). While this short-term exposure provides insight into immediate gene expression shifts, it may not fully reflect the physiological and molecular mechanisms that enable *P. vulnus* to persist under prolonged cold conditions (Towson and Lear, 1982, Acosta and Malek, 1979) (**Fig. 2A**). Principal component analysis (PCA) of the RNA-seq data confirmed high reproducibility among biological replicates and revealed transcriptional patterns associated with both developmental stages and environmental conditions (**Fig. 2B**, **2C**). For nematodes across different developmental stages, samples were clearly separated along principal component 1 (PC1), which explains 32.19% of the variance and broadly recapitulated progression through the nematode life cycle. PC2 (21.77%) distinguished egg samples from those of nematodes at post-hatching stages, likely reflecting gene expression patterns specific to early embryogenesis (**Fig. 2B**). PCA of RNA-seq data from nematodes experiencing different environmental conditions showed that PC1 (explaining 35.44% of the variance) strongly separated samples of cold-treated nematodes from ones of nematodes collected from grapevine and walnut hosts (**Fig. 2C**). Although this separation indicates a robust transcriptional response to cold exposure, some differences may reflect the fact that the cold-treated nematodes had originally been reared on grapevines. Additionally, PC2 clearly differentiated walnut-associated nematodes from grapevine-associated and cold-stressed individuals (**Fig. 2C**), indicating that the host environment also significantly influences the nematode gene expression.

**Fig. 2:**
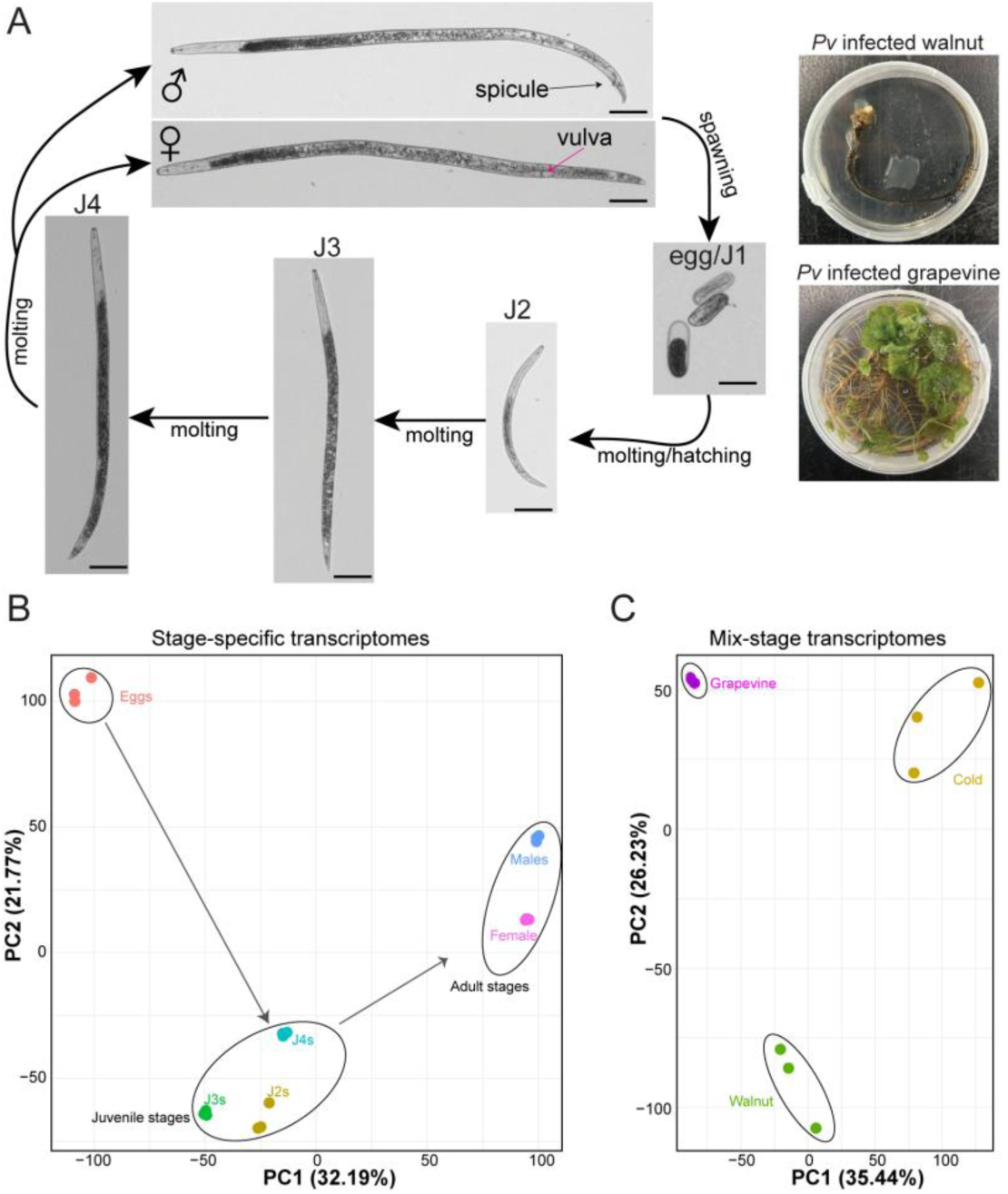
Gene expression dynamics in *Pratylenchus vulnus*. **A.** Representative micrographs of the six developmental stages of *P. vulnus*. Right: images of *P. vulnus*-infected walnut and grapevine roots. Scale bars: 50 µm. **B.** Principal component analysis (PCA) of RNA-seq data from biological replicates of individual developmental stages. Juvenile stages, adult stages, and eggs form distinct clusters. **C.** PCA of RNA-seq data from mixed-stage populations under three conditions: extracted from grapevine host, extracted from walnut host, and cold shock after extraction from grape. **B-C**. Each point represents a biological replicate.

To identify genes that significantly respond to developmental transitions or environmental conditions, we conducted a differential gene expression analysis. The largest transcriptional shifts during development—reflected by the greatest number of differentially expressed genes (DEGs; Padj < 0.05; log_2_ fold-change≥ 1)—occurred at two key transitions: from eggs to J2, and from J4 to adults (**Fig. 3A**). Notably, the egg-to-J2 transition involved substantial upregulation of genes encoding pectate lyases— enzymes involved in plant cell wall degradation—highlighting the activation of parasitism-related processes for host invasion and migration within the host (**Fig. 3B**). Additionally, genes encoding G protein-coupled receptors (GPCRs) were significantly enriched in J2 compared to eggs (**Fig. 3B and Fig. S3**). GPCRs constitute a large family of membrane receptors that mediate nematode responses to diverse external and internal signals, including small molecules and peptides (Cardoso et al., 2012, Beets et al., 2023, Holden-Dye and Walker, 2011). The pronounced stage-specific regulation of GPCR genes thus highlights their likely involvement in coordinating developmental processes and environmental responses during the lifecycle of *P. vulnus*.

**Fig. 3:**
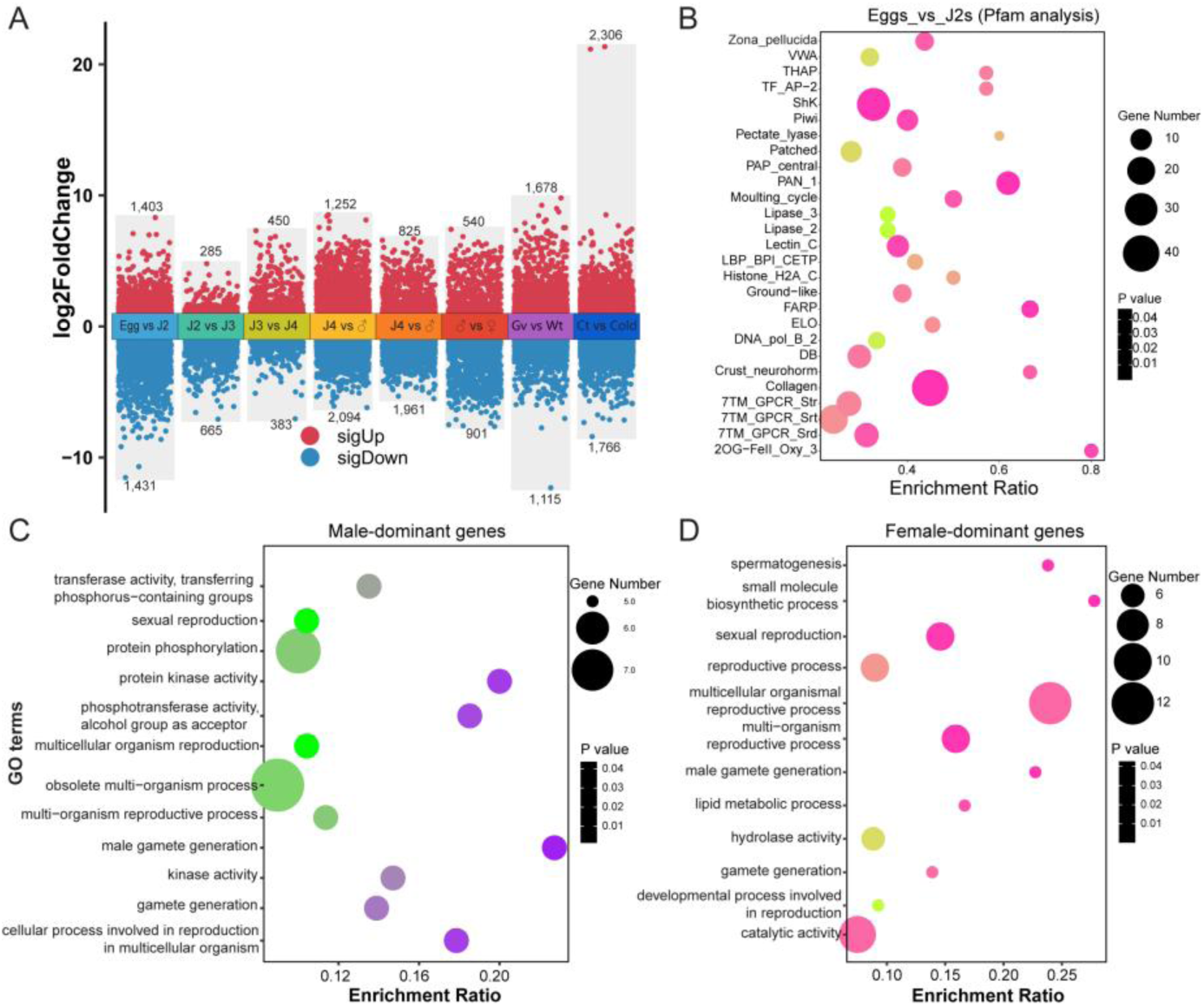
Differential gene expression and functional enrichment in *P. vulnus*. **A.** Differentially expressed genes (DEGs) between successive developmental stages, sexes, host conditions, and cold treatment in *P. vulnus*. Red and blue dots represent significantly upregulated (sigUp) and downregulated (sigDown) genes, respectively (‖log Foldchange‖ ≥ 1, adjusted p-value < 0.05). Numbers above and below each group indicate counts of up- and downregulated gene counts. Gv, grapevine; Wt, walnut; Ct, control. **B.** Pfam domain enrichment analysis of DEGs between egg and J2 juvenile stages. Dot size corresponds to the number of genes per Pfam domain; color intensity indicates significance (p-value). **C–D.** Gene Ontology (GO) enrichment analyses of male- and female-biased genes, respectively. Dot size corresponds to gene count; color reflects p-value.

Interestingly, the highest number of DEGs overall was observed after cold-treatment with 2,306 upregulated and 1,766 downregulated genes (**Fig. 3A**). Cold-responsive DEGs were primarily enriched for genes encoding transcription factors, ribosomal proteins, and regulators of cellular osmotic pressure (**Fig. S4**). When we compared the transcriptomes of *P. vulnus* parasitizing grapevine roots versus walnut roots, we detected 2,793 DEGs. Functional enrichment analysis of genes more highly expressed during walnut parasitism revealed significant enrichment in functions related to transcriptional regulation, DNA binding, morphogenesis, and tissue development, particularly epithelial tube (intestine) formation and organ morphogenesis (**Fig. S5**). Walnuts have a much tougher root than grapes. Therefore, this extensive transcriptional remodeling may facilitate development of stronger musculature, reinforced feeding structures, or enhanced absorptive tissues to exploit the physically demanding walnut root environment.

To characterize stage-specific gene expression dynamics in greater detail, we calculated transcripts per million (TPM) values for each gene and performed soft clustering analysis across all developmental stages. Each developmental stage exhibited a unique set (cluster) of highly expressed genes (**Figs. S6 and S7**). Functional enrichment analyses of these clusters revealed stage-specific biological functions. For instance, clusters with male-biased expression were enriched for terms related to male gamete generation, reproduction, and kinase activity (**Fig. 3C**). Gene clusters with female-biased expression were enriched for reproduction-related functions, lipid metabolism, and small-molecule metabolism (**Fig. 3D**). Interestingly, genes that were highly expressed in females also exhibited enrichment for the GO terms male gamete generation and spermatogenesis, suggesting that some sex-associated genes have biological functions beyond sex determination alone.

Finally, clustering analysis incorporating cold-induced RNA-seq data identified a subset of genes that are specifically and strongly expressed under cold conditions (cold-induced cluster; **Fig. S8A**). Functional enrichment analysis of these cold-responsive genes revealed an enrichment of transcription factors, particularly members of the basic leucine zipper (bZIP) family (**Fig. S8B**), a protein family previously associated with responses to cold in plants (Han et al., 2020, Liu et al., 2019, Liu et al., 2023, Kreps et al., 2002). Genes encoding Na⁺/H⁺ exchangers—known regulators of osmotic balance—were also significantly enriched among cold-responsive genes in Drosophila (Enriquez and Colinet, 2019). Although these results highlight the nimble transcriptional responses of *P. vulnus* under short-term cold exposure, additional long-term studies are needed to capture the full scope of its cold-acclimation strategies. Collectively, these transcriptomic data highlight the dynamic and context-dependent nature of gene expression throughout *P. vulnus* development and parasitism, providing valuable insights into its development and adaptation to specific hosts and environmental conditions.

### Expression landscape and genomic distribution of effector and HGT-derived genes

PPNs use effector proteins to facilitate infection and suppress defenses in host plants (Kud et al., 2019). To better understand the genomic features associated with adaptation to a parasitic lifestyle, we examined the distribution and expression patterns of secreted proteins as most characterized effectors of PPNs are secreted (Rehman et al., 2016). We identified 1,629 candidate effector genes, which were relatively evenly distributed across all chromosomes (**Fig. 4A**). To characterize the developmental regulation of these effector candidates, we calculated the TPM values of each gene across nematode development and performed clustering analysis. Although *P. vulnus* is a migratory PPN capable of infecting plants continuously from the J2 stage onward, these candidate effector genes exhibited distinct and surprisingly stage-specific expression profiles (**Fig. 4B and Fig. S9**). Such stage-specific expression patterns likely reflect diverse functional demands at each developmental stage—such as root penetration, immune evasion, and nutrient acquisition. In addition, all nematodes, both free living and parasitic, produce a repertoire of secreted proteins that function in their development, neurotransmission and other functions. Thus, while secreted proteins are likely enriched in effectors, some of the candidate effectors may, in fact, have other functions. One clear example is a cluster dominated by genes expressed almost exclusively in eggs, which are most likely secretory proteins with developmental rather than parasitic roles.

**Fig. 4:**
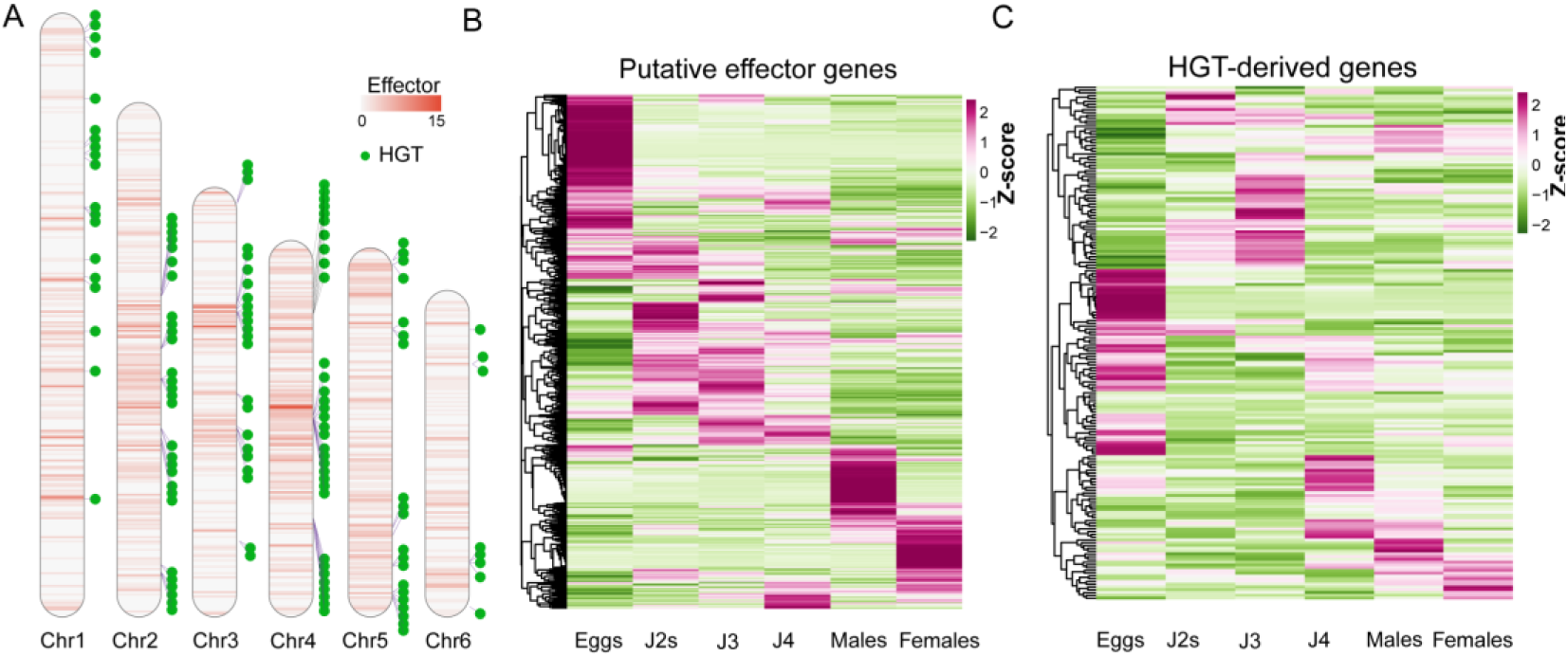
Genomic distribution and developmental expression of effector and horizontally transferred (HGT) genes in *Pratylenchus vulnus*. A. Chromosomal distribution of putative effector genes (heatmap) and horizontally transferred genes (green dots) across the six *P. vulnus* chromosomes. Effector gene density is shown as a heatmap (0–15 per window), while HGT-derived gene positions are marked as individual points. **B**. Heatmap showing expression (Z-score normalized TPM) of putative effector genes across developmental stages. **C**. Heatmap of expression patterns for HGT-derived genes across the same developmental stages.

PPNs carry an array of genes that have signatures of having been acquired by HGT involved in carbohydrate metabolism, many of which have been demonstrated to have important functions in parasitism (Mitreva et al., 2009). In contrast to the dispersed genomic arrangement of secreted-protein genes, HGT-derived genes exhibited distinctly clustered genomic distributions (**Fig. 4A**). This clustered arrangement aligns with previous observations in RKNs (Dai et al., 2023), suggesting that these genes may have expanded in copy number following acquisition. Similar to secreted-protein genes, HGT-derived genes displayed clear stage-specific expression patterns, with distinct subsets highly expressed at specific developmental stages (**Fig. 4C**).

Functional enrichment analysis of HGT-derived genes identified a significant representation of glycosyl hydrolases and pectate lyases (**Fig. S10**); these effectors are essential for plant cell wall degradation and sugar metabolism during infection (Rai et al., 2015, Dayi, 2024). All identified pectate lyase genes in *P. vulnus* appear to be derived from HGT events. Together, these findings reveal distinct genomic distributions and precisely coordinated expression patterns of secreted protein and HGT-derived genes, offering valuable insights into the biological adaptations to parasitic lifestyle of *P. vulnus*.

### Genome comparison with other nematode species reveals extensive chromosomal rearrangements and unique genomic regions in *P. vulnus*

To better understand genomic adaptations associated with plant parasitism in *P. vulnus*, we conducted a comparative genomic analysis that included several closely related PPNs with sequenced genomes. A recent BUSCO50-based phylogeny identified the migratory PPN genus *Pratylenchus* as most closely related to the sedentary PPN genus *Meloidogyne* (Qing et al., 2024). Thus, we first evaluated the chromosomal collinearity between *P. vulnus* and *M. graminicola*, a well-characterized diploid root-knot nematode with a similar genome size, but 18 chromosomes **(Fig. 5A)**. This comparison revealed blocks of synteny, but also several examples where individual chromosomes in *P. vulnus* exhibited collinearity to multiple chromosomes in *M. graminicola*. Notably, chromosome 3 of *P. vulnus* contained collinear segments corresponding to eight separate chromosomes in *M. graminicola* (**Fig. 5A**). Conversely, several chromosomes in *M. graminicola* each had segments represented on only single chromosomes in *P. vulnus* (**Fig. 5A**).

**Fig. 5:**
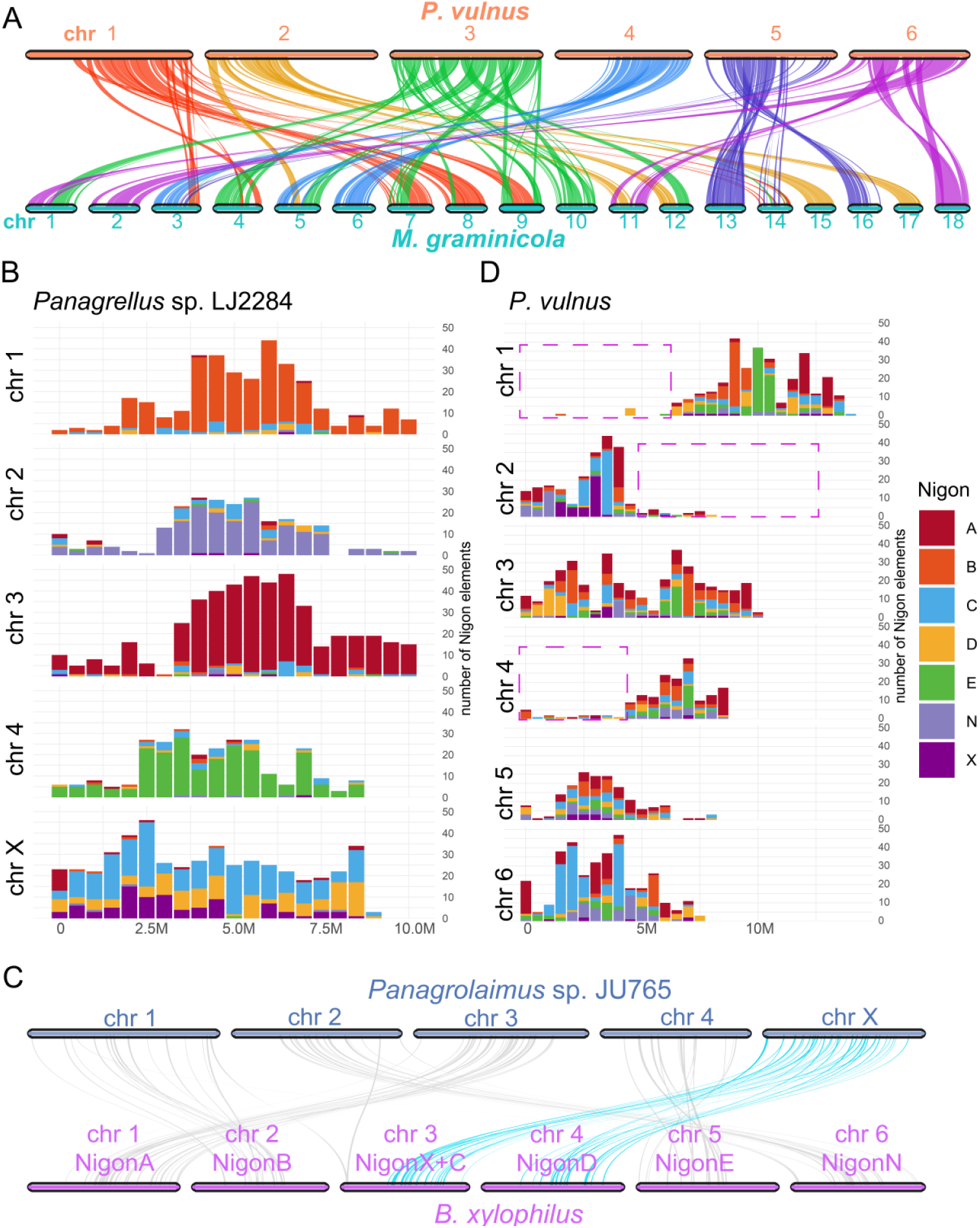
Chromosomal collinearity and Nigon element distribution in *Pratylenchus vulnus* and related nematodes. **A.** Chromosomal collinearity between *P. vulnus* and *Meloidogyne graminicola*. Colored ribbons link orthologous genes, with colors corresponding to their P*. vulnus* chromosome of origin. **B.** Distribution of Nigon elements across chromosomes in *Panagrellus* sp. LJ2284. Most autosomes are dominated by a single Nigon element, while the X chromosome contains a mosaic of three. **C.** Syntenic relationships between the X chromosome of *Panagrellus* sp. JU765 and chromosomes 3 and 4 of *Bursaphelenchus xylophilus*. **D.** Distribution of Nigon elements across *P. vulnus* chromosomes. Each bar represents a genomic window and is color-coded by Nigon element identity. Regions without detectable Nigon assignment are highlighted with dashed purple boxes.

Intriguingly, the left arms of chromosomes 1 and 4 as well as the right arm of chromosome 2 in *P. vulnus* lacked corresponding sequences in the *M. graminicola* genome (**Fig. 5A**), suggesting that these genomic regions were acquired in *P. vulnus* after splitting from common ancestor with Mg or were lost in Mg. Interestingly, these regions correspond roughly to those noted to have low %GC and low gene density **(Fig. 1D)**. Further comparative analyses with other migratory but more distantly related PPNs—*Ditylenchus destructor*, *Aphelenchoides besseyi*, and *Bursaphelenchus xylophilus*—as well as relatively closely related free-living nematodes of the latter two species—*Panagrellus* sp. LJ2284 (**Fig. S11**) and *Panagrolaimus* sp. JU765—similarly failed to identify homologous genomic segments, highlighting their uniqueness (**Fig. S12A–D**).

We extended our comparative analysis to include the published genome of *P. scribneri* (Arora et al., 2023), a congener of *P. vulnus*, to further investigate the origin of the non-collinear chromosomal regions. Although the *P. scribneri* genome is somewhat fragmented, we identified partial homology between its sequences and the left arms of chromosomes 1 and 4 as well as the right arm of chromosome 2 in *P. vulnus*—regions that otherwise showed poor synteny with *M. graminicola* and other nematode genomes **(Fig. S13)**. This suggests that these segments may be partially conserved within the genus *Pratylenchus* and may trace back to an ancestral lineage common to this genus.

Nigon elements, which are conserved genomic blocks present across Rhabditida nematodes, serve as useful markers for analyzing chromosome evolution due to their characteristic distribution patterns among nematode chromosomes (Gonzalez de la Rosa et al., 2021). Analysis of Nigon element distribution in *Panagrolaimus* sp. JU765 and *Panagrellus* sp. LJ2284 showed that each autosome primarily contained a single Nigon element, while their X chromosomes originated from ancestral chromosome block fusions involving NigonX, NigonC, and NigonD elements (**Fig. 5B and Fig. S14**). Similarly, the six chromosomes of *B. xylophilus* predominantly contained single Nigon elements, except for its X chromosome, which arose through the fusion of ancestral NigonX and NigonC chromosomal segments (Gonzalez de la Rosa et al., 2021). Furthermore, collinearity analysis demonstrated that the X chromosome of *Panagrolaimus* sp. JU765 shared collinear segments with chromosomes 3 (NigonX, NigonC) and 4 (NigonD) of *B. xylophilus* (**Fig. 5C**), reinforcing the utility of Nigon elements for inferring the evolutionary histories of chromosomes. In comparison, the distribution of Nigon elements in *P. vulnus* chromosomes showed a notably fragmented and complex pattern (**Fig. 5D**), indicating extensive rearrangements, fusions, or fragmentation events compared to ancestral chromosome structures. Integrating Nigon element (NigonX) distribution data with interspecies protein-based synteny analyses (**Fig. 5D and Fig. S12**), we propose that chromosome 2 is likely the X chromosome of *P. vulnus*.

Remarkably, the left arms of chromosomes 1 and 4, as well as the right arm of chromosome 2 in *P. vulnus* were nearly devoid of Nigon elements. Given the lack of protein-coding gene collinearity between these regions and other examined nematodes **(Fig. 5A and Fig. S12**), we speculate that these segments may have originated from nematodes lacking typical Nigon elements or even other phyla. Collectively, our results demonstrate that the genome of *P. vulnus* has undergone significant chromosomal rearrangements and genomic innovations compared to its closest relatives. These historical events may have shaped its ecological specialization on woody plants by incorporating genomic elements from diverse ancestral sources, underscoring a complex evolutionary trajectory that is distinct from that of other related nematodes.

### Comparative analysis reveals lineage-specific diversification of cell wall– degrading enzymes in *P. vulnus* and related nematodes

*P. vulnus* is one of the most damaging PPNs of woody host plants such as walnut and peach (*Prunus persica*), which have highly lignified and structurally complex cell walls. This specialized ecological niche raises the question of whether *P. vulnus* possesses a distinct repertoire of cell wall degrading enzymes (CWDEs) that enable it to colonize such robust tissues. To address this, we performed a comparative analysis of CWDE gene families across eight nematode species spanning a spectrum of lifestyles, from free-living to highly specialized plant parasites (Qing et al., 2024). These included the free-living species *Oscheius tipulae* (Ot), *Panagrolaimus* sp. JU765, and *Panagrellus* sp. LJ2284; the migratory parasites *B. xylophilus* (Bx), *A. besseyi* (Ab), and *D. destructor* (Dd), and *P. vulnus* (Pv); and the sedentary endoparasite *M. graminicola* (Mg). We focused on CWDE families including glycoside hydrolases (GH), pectate lyases (PL) and carbohydrate-binding modules (CBM13) (Rai et al., 2015, Dayi, 2024, Gahoi and Gautam, 2017). Among the analyzed nematodes, *B. xylophilus* had the largest CWDE gene complement (152 genes), followed by the free-living species *Panagrolaimus* sp. JU765 (101 genes) and *Panagrellus* sp. LJ2284 (78 genes) (**Fig. 6A and Table S2**). By contrast, the sedentary parasite *M. graminicola* possessed the smallest CWDE repertoire, comprising only 28 genes. *P. vulnus* exhibited an intermediate-sized CWDE gene set (66 genes), placing it between free-living nematodes and sedentary plant parasites (**Fig. 6A and Table S2).** Interestingly, although both *P. vulnus* and *B. xylophilus* parasitize woody hosts, their CWDE gene compositions differed markedly (**Table S2**).

**Fig. 6:**
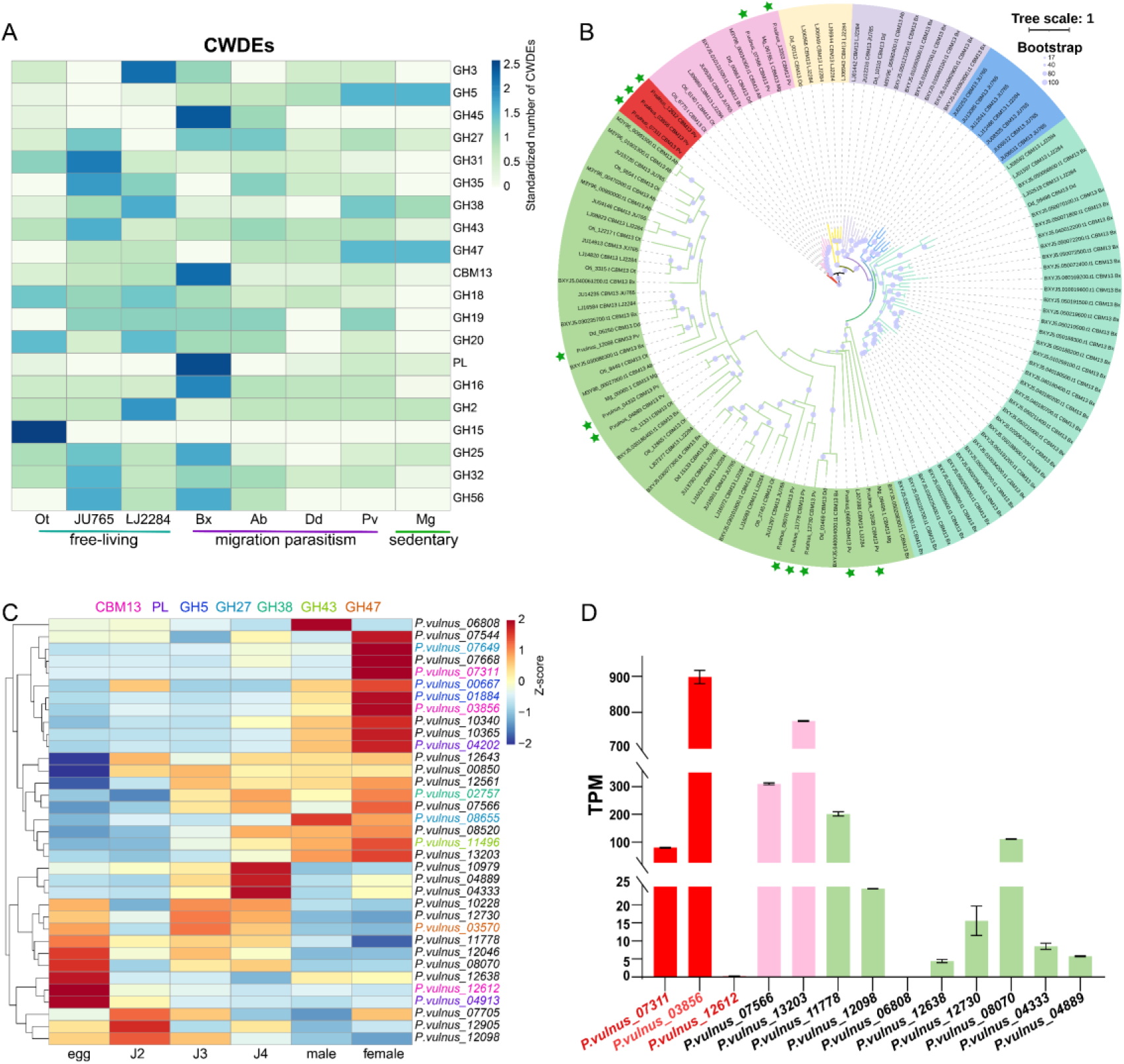
Characterization of cell wall degrading enzymes (CWDEs) across nematode species. **A.** Abundance of CWDE gene families across eight nematode species, normalized by min–max scaling. Species are grouped by ecological lifestyle: Free-living: *Oscheius tipulae* (Ot), *Panagrolaimus* sp. JU765, *Panagrellus* sp. LJ2284. Migratory plant parasites: *Bursaphelenchus xylophilus* (Bx), *Aphelenchoides besseyi* (Ab), *Ditylenchus dipsaci* (Dd), *Pratylenchus vulnus* (Pv). Sedentary endoparasite: *Meloidogyne graminicola* (Mg). Gene families include glycoside hydrolases (GHs), pectate lyases (PL), and carbohydrate-binding modules (CBMs). **B.** Phylogenetic tree of CBM13 proteins from all eight nematode species. Colored branches represent distinct clades. Asterisks indicate *P. vulnus* genes that form phylogenetically distinct lineages, lacking close homologs in other nematodes. Tree constructed using IQ-TREE with 1,000 ultrafast bootstrap replicates. **C.** Heatmap showing stage-specific expression (Z-score of TPM) for various *P. vulnus* CWDE gene families. Gene names are colored according to gene family: CBM13 (magenta), PL (purple), GH5 (blue), GH27 (cyan), GH38 (green), GH43 (lime green), GH47 (orange). Colored gene names indicate *P. vulnus*–specific genes that form phylogenetically distinct clades (i.e., without homologs in other nematodes), while black-labeled genes are conserved and cluster with homologs from other species.**D.** RNA-seq–based expression of CBM13 genes in *P. vulnus* females. Colors match clades from panel B.

To explore this further, we conducted phylogenetic analyses of key CWDE families enriched in *P. vulnus*, including GH5, GH27, GH38, GH43, GH47, CBM13, and PL. These analyses indicated that several *P. vulnus* CWDEs formed clades that were phylogenetically distinct from those in other nematodes, suggesting lineage-specific expansions or independent origins (**Fig. 6B and Fig. S15, 16)**. For instance, the CBM13 family, associated with xylan degradation (Chen et al., 2020), is abundant across nematodes, particularly in *B. xylophilus* (48 genes) and free-living species such as *Panagrellus* sp. LJ2284 (18 genes). Phylogenetic analysis grouped CBM13 proteins into seven distinct clades (**Fig. 6B**). While some CBM13 genes in *P. vulnus* grouped closely with homologs from other nematodes, three CBM13 genes formed a distinct lineage separate from that of other nematodes, indicating a possible independent evolutionary origin or diversification. Similar patterns were observed for other CWDE families, including GH5 (7 genes), GH38 (4 genes), GH43 (3 genes), and PL (5 genes), which had expanded in *P. vulnus* relative to *M. graminicola* (**Table S2 and Fig. S15, 16**).

Further analysis revealed that many of the phylogenetically distinct CWDE genes in *P. vulnus* are located in non-syntenic regions on chromosomes 1, 2, 4, and 5 (**Fig. S17**), suggesting possible origins through lineage-specific innovation or horizontal gene transfer. To investigate their functional relevance, we examined developmental stage–specific expression patterns for various CWDE gene families in *P. vulnus*. While no consistent pattern emerged at the level of gene families, several *P. vulnus*–specific CWDE genes showed elevated expression during parasitic stages (J2– J4) and in adult females (**Fig. 6C**). This includes the CBM13 gene *P. vulnus_03856*, which exhibited the highest expression among all CWDEs in females (**Fig. 6D**). Although the biases in expression patterns across developmental stages were gene-specific rather than family-wide, the transcriptional activity of several phylogenetically distinct genes during parasitic stages underscores their potential relevance to host invasion and colonization. Together, these findings suggest that *P. vulnus* has evolved a unique set of CWDEs through lineage-specific diversification and expression, likely enabling adaptation to the physical and chemical defenses of woody plant tissues.

## Discussion

In this study, we present a chromosome-scale genome assembly for *P. vulnus*, consisting of 6 chromosomes corresponding to the previously reported chromosome number for this strain. One of the most striking findings was the presence of extensive genomic regions that lack synteny with other sequenced nematode genomes, including well-characterized free-living and parasitic species. These non-collinear regions were structurally distinct and compositionally divergent, exhibiting GC content as low as 25– 30%, in contrast to the 32–55% GC content observed in collinear, conserved chromosomal segments. The presence of these low-GC, non-syntenic regions suggests a possible origin from evolutionarily distant nematode lineages, potentially from other clades or orders for which genomic resources are still unavailable. This hypothesis is supported by the distribution of parasitism-related genes in these regions. Several genes encoding cell wall–degrading enzymes (CWDEs), such as those from the CBM13, GH5, GH27, GH38, GH43, and PL families, are in these non-syntenic blocks and show distant phylogenetic relationships to homologs in other nematodes. Likewise, a subset of predicted effector genes—key to host manipulation—are found within these regions, several of which exhibit stage-specific expression across juvenile and adult phases. Alternatively, these regions may represent ancient HGT events or remnants of genome fusion, possibly derived from now-extinct or unsampled species. Their divergence in base composition and lack of conserved gene order support the hypothesis that they evolved under distinct selective pressures or genomic contexts, separate from the core genome.

In addition to shedding light on genomic novelty within *P. vulnus*, our findings have broader implications for the evolution of PPNs karyotypes. Notably, the observed absence of collinearity in certain chromosomal regions parallels chromosomal plasticity observed in other parthenogenetic nematodes, particularly RKNs. These nematodes also display extraordinary variation in chromosome number (Roman and Triantaphyllou, 1969), extensive structural rearrangements, and evidence of ancient hybridization and polyploidy. The fact that several *Pratylenchus* species reproduce by mitotic parthenogenesis, with chromosome numbers ranging from 5 to 32, suggests that *Pratylenchus* may represent a pivotal lineage in understanding the evolutionary origin of the highly derived genomes of *Meloidogyne*. Intriguingly, the lowest chromosome number reported in *Meloidogyne* is 4 (Alvarez-Ortega et al., 2019), similar to that observed in some *Pratylenchus* species. This small chromosome count raises the possibility that extensive chromosomal rearrangements and reductions may have accompanied or facilitated transitions in reproductive mode and host adaptation. The *Meloidogyne* genomes sequenced so far lack telomeres, whereas *P. vulnus* has long runs of canonical telomere repeats.

To test these evolutionary scenarios more robustly, additional high-quality genome assemblies from other *Pratylenchus* and *Meloidogyne* species are needed. In particular, sequencing *Pratylenchus* species with extreme karyotypes (Roman and Triantaphyllou, 1969) or unique ecological niches could help reconstruct ancestral genomic states and delineate the steps underlying chromosome reshaping, gene acquisition, and reproductive transitions in PPNs.

In conclusion, our study highlights the evolutionary complexity and genomic plasticity of *P. vulnus*, a key representative of woody plant–parasitizing nematodes. The discovery of non-collinear, low-GC chromosomal regions points to deep evolutionary events that shaped this genome, potentially involving ancient lineage divergence or acquisition from uncharacterized taxa. Moreover, our findings underscore the importance of expanded taxon sampling and chromosome-resolved assemblies in uncovering the origins of genomic novelty and adaptation in parasitic nematodes.

## Methods

### Nematode culture and collection

*Pratylenchus vulnus* nematodes were propagated on sterile grape and walnut roots grown on solidified Driver Kuniyuki Walnut (DKW) basal medium (i.e. no growth regulators) under axenic conditions. After 3 weeks of root growth, sterile nematodes were introduced, and the cultures were maintained at 25°C for 6–8 weeks. For nematode collection, plant roots were finely chopped with a scalpel along with the surrounding medium and transferred to Baermann funnels lined with a single layer of Kimwipes. Water was added until the plant material was fully submerged, and nematodes were allowed to migrate for approximately 4 h. The eluate was then collected from the bottom of the funnel and mixed with 70% sucrose to achieve a final concentration of 35%. The mixture was transferred to a 1.5 mL microcentrifuge tube and centrifuged at 5,000 rpm for 2 min. The pellet, consisting of impurities, was discarded. The supernatant containing the nematodes was then passed through a 3 μm filter membrane to remove the sucrose, yielding purified nematodes. The purified nematodes were used in all experiments. Genomic DNA used for sequencing was extracted from *P. vulnus* infecting grape roots. Transcriptome data from different developmental stages and cold treatment conditions were obtained from nematodes infecting grapevine roots; however, the mixed-stage transcriptome samples were collected from nematodes infecting both grape and walnut roots.

### Acquisition of nematodes at different stages of development and under cold stress

To collect nematodes at different stages of development, individuals were manually picked under a stereomicroscope. Eggs were the most easily distinguishable stage, and approximately 150 eggs were collected for each biological replicate. Newly hatched juveniles from eggs were designated as second-stage juveniles (J2s), with ∼150 individuals collected per replicate. Third-stage (J3) and fourth-stage (J4) juveniles were selected based on body size differences from a mixed-stage nematode population, with ∼100 individuals per replicate. Adult females and males were distinguished by their reproductive organs, and ∼50 individuals of each sex were collected per replicate.

### DNA and RNA extraction and library construction

DNA and RNA were extracted from *P. vulnus* using previously established protocols for root-knot nematodes(Dai et al., 2023). Briefly, for DNA extraction, mixed-stage nematodes were suspended in 200 µl lysis buffer, flash-frozen in liquid nitrogen, and ground into a fine powder using a mortar and pestle. After adding another 300 µl lysis buffer, the lysate was mixed with 500 µl CTAB buffer at 65°C for 30 min, followed by protein removal using phenol:chloroform:isoamyl alcohol extraction. The DNA was precipitated with ethanol. DNA for Illumina sequencing was used directly after extraction, and high-molecular-weight DNA was further purified for Nanopore and PacBio HiFi sequencing using magnetic beads before being submitted to Novogene (Sacramento, CA) for library preparation and sequencing. Hi-C library construction was performed by Phase Genomics using mixed-stage nematodes. Total RNA was extracted from the samples using RNeasy Mini Kit (QIAGEN, 74106), and RNA-seq libraries were constructed using a VAHTS Universal V10 RNA-seq Library Prep Kit for Illumina (Vazyme, NR606-01). Libraries were sequenced on an Illumina NovaSeq instrument as 150-bp paired-end reads by NOVOGENE.

### Assessment of genome size and ploidy

The size, heterozygosity, and ploidy level of the *P. vulnus* genome were estimated based on Illumina short-read sequencing data. *K*-mer-based ploidy analysis was performed using Smudgeplot (Ranallo-Benavidez et al., 2020) to confirm that *P. vulnus* is a diploid organism. Genome size and heterozygosity were estimated using GenomeScope2 (Ranallo-Benavidez et al., 2020) following methods that were previously applied to root-knot nematodes (Dai et al., 2023).

### Contig assembly

The *P. vulnus* genome was first assembled using long-read Nanopore sequencing data with default parameters in the NECAT (Chen et al., 2021) assembler. This preliminary assembly was then error-corrected and gap-filled using PacBio HiFi reads. Subsequently, three rounds of polishing were performed with Illumina short-read data using Pilon (Walker et al., 2014) to improve sequence accuracy. The error-corrected genome assembly was used for chromosome anchoring.

### Chromosome-level genome construction and identification of telomeric elements

The *P. vulnus* contigs were anchored onto chromosomes using Hi-C data, which captured chromatin interaction information. First, the Hi-C reads were aligned to the contig-level assembly using the BWA (v0.7.17-r1188) (Li and Durbin, 2009) and Juicer (v1.5.7) pipelines (Durand et al., 2016b). The 3D-DNA (v180419) pipeline (Dudchenko et al., 2017) was then employed to scaffold adjacent genomic fragments based on chromatin contact signals, generating a draft chromosome-scale assembly. The resulting “.hic” and “.assembly” files were manually curated and corrected using Juicebox (Durand et al., 2016a) to produce the final chromosome-level genome assembly. An in-house Python script was used to identify the tandemly repeated telomeric repeat sequence “TTAGGC”.

For *Panagrellus* sp. LJ2284, the contig assembly was downloaded from NCBI under accession number GCA_024447225.1, along with the corresponding Hi-C dataset (accession number SRR23107840 from the Sequence Read Archive [SRA] at NCBI). A chromosome-level genome was assembled using the strategy described above. During assembly, redundant “unzipped” contigs were identified and removed, likely representing alternative haplotypes. The final assembly consisted of five chromosomes, with a total genome size of 65.4 Mb. This chromosome-level *Panagrellus* sp. LJ2284 genome assembly was used for genome annotation.

### Genome annotation

To annotate protein-coding genes in *P. vulnus*, transcriptome data were integrated from all developmental stages to perform comprehensive gene predictions. First, repetitive sequences in the genome were masked using RepeatMasker (Tarailo-Graovac and Chen, 2009). Subsequently, transcriptome data from eggs, J2s, J3s, J4s, males, females, and mixed samples were sequentially aligned to the genome using HISAT2 (Kim et al., 2019). Low-quality reads (mapping quality [MAPQ] < 30) were removed using SAMtools (Li et al., 2009) to generate the “final.bam” files, which were used for multiple gene predictions. The alignments were used as input for BRAKER3 (Gabriel et al., 2024) to perform *ab initio* gene predictions based on GeneMark-ET/EP and AUGUSTUS, generating the file “braker.gff3”. In parallel, TransDecoder (https://github.com/TransDecoder/TransDecoder) and StringTie (Pertea et al., 2015) were used on the same .bam files to predict protein-coding genes, resulting in the file “stringtie.gff3”. Additionally, all RNA-seq datasets were assembled *de novo* using Trinity (Grabherr et al., 2011), and the assembled transcripts (Trinity.fasta) were used as input for PASA (Haas et al., 2008) to identify gene structures, producing the file “pasa_assemblies.gff3”. Finally, the three annotation sources were combined using EVidenceModeler (Haas et al., 2008), with weights set to 4 for AUGUSTUS, 4 for TransDecoder, and 5 for PASA. The merged annotation file (EVM.merged.gff3) was processed with the gffrename.py script to standardize gene names and obtain the final gene annotation file. The Launch_PASA_pipeline.pl was used to annotate isoforms.

For genome annotation of *Panagrellus* sp. LJ2284 and *Panagrolaimus* sp. JU765 (GCA_964245515.1), RNA-seq data were downloaded from the SRA database under accession numbers SRR22163891 and SRR5253561, respectively. The two genomes were annotated using the strategy described above, yielding 16,973 and 16,343 protein-coding genes, respectively. These annotated genomes were used for subsequent analysis. The predicted genes were annotated with GO and KEGG terms using eggNOG-mapper (Cantalapiedra et al., 2021), and Pfam domains were identified using the Pfam_scan.pl script (Mistry et al., 2021). GO enrichment analysis was performed using TBtools-II (Chen et al., 2023), while Pfam enrichment analysis was conducted with the Pfam_Enrichment.py script.

### Predicting transposable elements, horizontal gene transfer events, transcription factors, and secreted proteins

The EDTA (v2.2.0) pipeline (Ou et al., 2019) was employed to predict TEs in the *P. vulnus* genome, with the parameters “--overwrite 1 --force 1 --sensitive 1 --anno 1 -- evaluate 1”. Horizontal gene transfer events were predicted using the AvP pipeline (Koutsovoulos et al., 2022) with the UniRef90 database. Transcription factors were identified using the AnimalTFDB (v4.0) database (Shen et al., 2023). Secreted proteins were predicted by first identifying signal peptides using SignalP5 (Almagro Armenteros et al., 2019), followed by the detection of transmembrane domains using TMHMM (Krogh et al., 2001). Proteins with signal peptides but lacking transmembrane domains were classified as putative secreted proteins.

### Analysis of genome collinearity between different nematodes

Due to the distant evolutionary relationships among the nematode species, inter-genomic synteny analysis was conducted based on annotated protein sequences to minimize the impact of synonymous mutations. Homologous proteins between each pair of genomes were identified using MCScan (Wang et al., 2012). First, the gene list in BED format was generated using the jcvi.formats.gff script, and all protein sequences from each genome were extracted using gffread (Pertea and Pertea, 2020). Cross-species homologous protein sequences were identified using jcvi.compara.catalog, and a synteny analysis of the homologs was performed with jcvi.compara.synteny using the parameter --minspan=10. Finally, interspecies chromosomal synteny was visualized using jcvi.graphics.karyotype.

### RNA-seq and expression profile analyses

For differential gene expression analysis, the transcriptome data were aligned to the genome assembly generated in this study using HISAT2. The gene-level read counts were then quantified using HTSeq-count (v0.13.5) (Anders et al., 2015). Significance testing for DEGs was performed using DESeq2 (Love et al., 2014): genes with an adjusted *p*-value (Padj) < 0.05 and an absolute log_2_(fold-change) > 1 were considered to be DEGs. To calculate gene expression values as TPM counts for each developmental stage, STAR was used (Dobin et al., 2013) for alignment and RSEM (Li and Dewey, 2011) for quantification. Expression profiles was conducted using Mfuzz (Kumar and Futschik, 2007), and heatmaps were visualized using the pheatmap R package.

### Phylogenetic analysis of cell wall–degrading enzymes

DIAMOND (Buchfink et al., 2015) was used to align nematode protein sequences against the “CAZy.dmnd” database to identify CAZy family proteins. Based on previous reports (Rai et al., 2015, Gahoi and Gautam, 2017), CWDEs from each nematode species were extracted according to their respective CAZy family classifications. These sequences were aligned using MAFFT (Katoh and Standley, 2013), and poorly aligned regions were trimmed using trimAl (Capella-Gutiérrez et al., 2009). Phylogenetic trees were reconstructed using IQ-TREE (Nguyen et al., 2015) and visualized with iTOL (Letunic and Bork, 2024).

## Availability of data and materials

Requests for further information and resources should be directed to the lead contact, Shahid Siddique. This study did not generate any new reagents. All sequencing data have been deposited in the Sequence Read Archive (SRA) at the National Center for Biotechnology Information (NCBI). The Nanopore, HiFi, Illumina, Hi-C, and RNA-seq raw data of *P. vulnus* were deposited under BioProject PRJNA1283215 (31 SRA files); the genomes of *Panagrellus* sp. LJ2284 and *Panagrolaimus* sp. JU765 were downloaded from NCBI under accession numbers GCA_024447225.1 and GCA_964245525.1, respectively. The RNA-seq data from *Panagrellus* sp. LJ2284 and *Panagrolaimus* sp. JU765 were downloaded from the SRA under accession numbers SRR22163891 and SRR5253561, respectively. The Hi-C data for *Panagrellus* sp. LJ2284 were downloaded from the SRA under accession number SRR23107840.

## Acknowledgments

We thank the University of California Davis farm server system for their support with data analysis.

## Author contributions

S.S. conceived and administered the project and acquired the funding; D.D. performed the data analysis; D.D. and Y.Z. generated the figures; D.D., Y.Z., and R.L. collected the stage-specific nematodes; D.D. constructed the stage-specific RNA-seq libraries; X.Y. and B.C. raised the nematodes; D.D. wrote the draft manuscript; S. C. G and S.S. revised the manuscript. All authors contributed to the manuscript.

