## Supplementary Figure1 for "Genomic and Transcriptomic Insights into the Evolution and Parasitic Strategy of the Woody-Plant Nematode *Pratylenchus vulnus*"

**Supplementary Figures:**

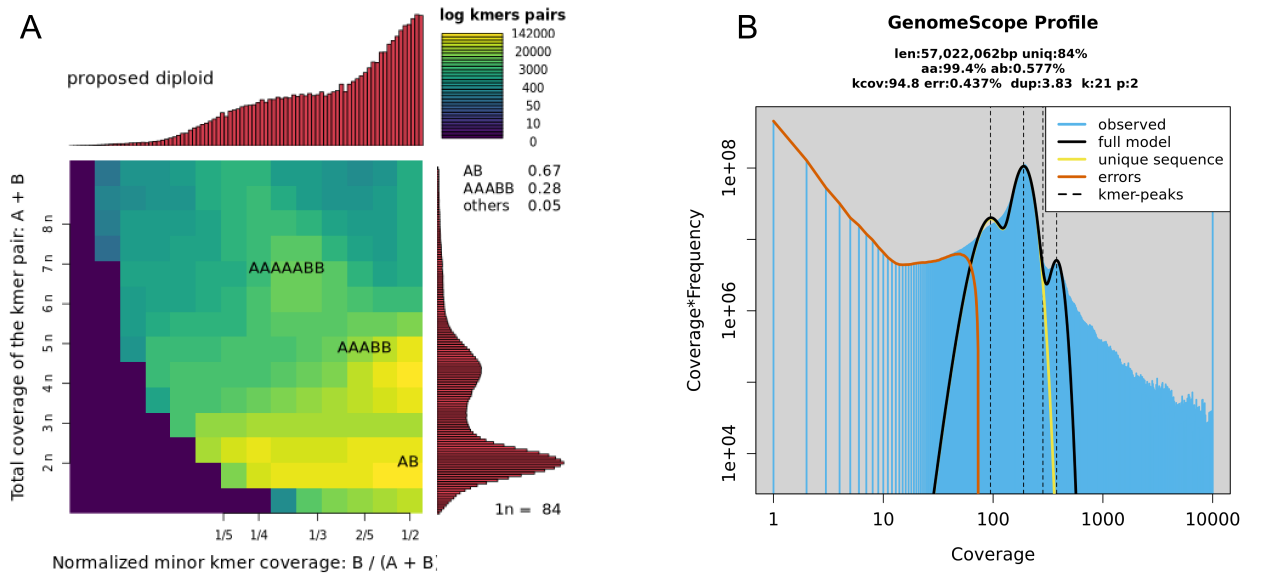

**Supplementary Fig. 1: Genome ploidy and size estimation of *P. vulnus*.** (**A**) Smudgeplot analysis indicates that the genome of *P. vulnus* is diploid. The primary peak corresponding to AB k-mer pairs supports a diploid genome structure. (**B**) GenomeScope2 profile estimating the genome size of P. vulnus at approximately 57.02 Mb, with high k-mer coverage (kcov = 94.8), low error rate (err = 0.437%), and moderate duplication rate (dup = 3.83%).

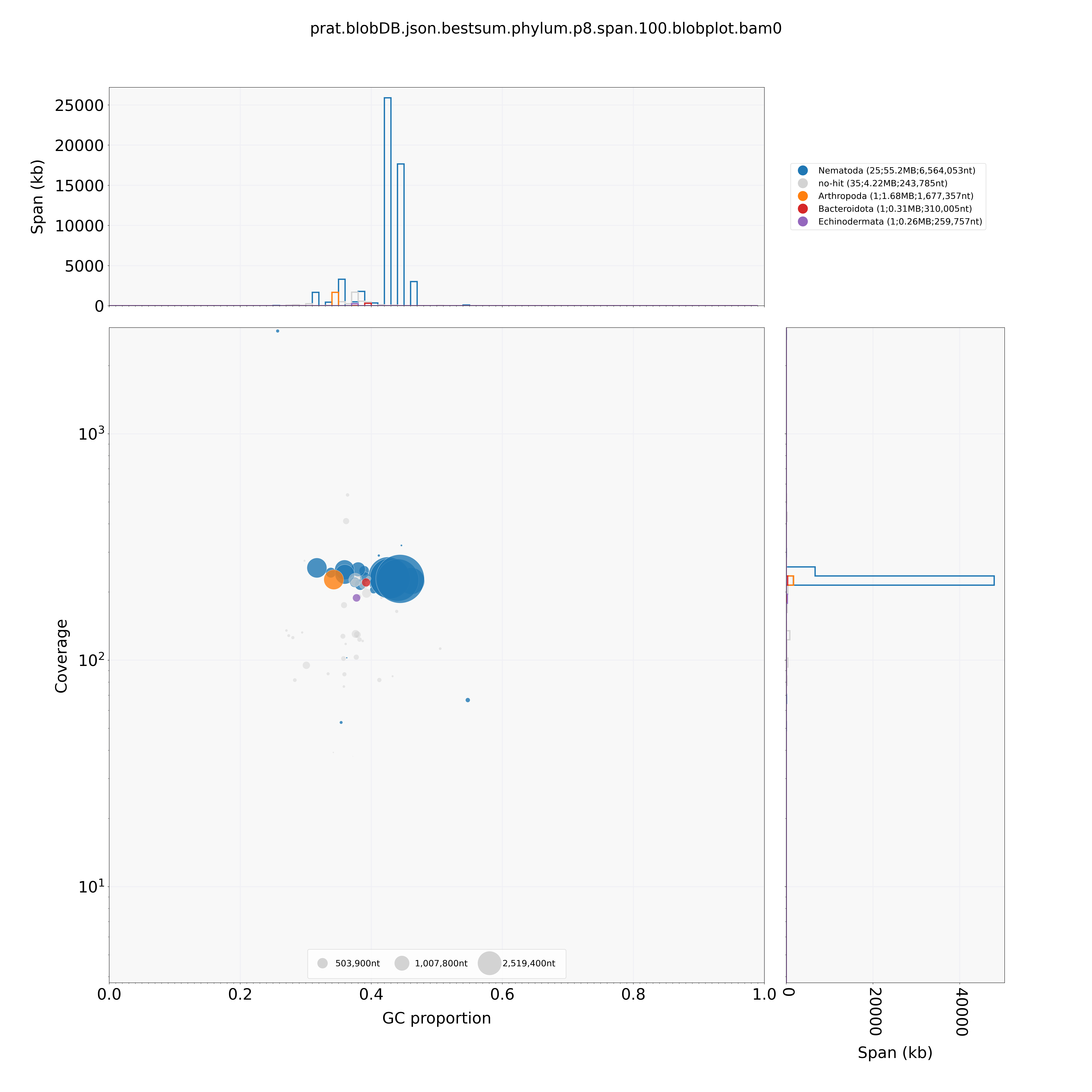

**Supplementary Fig. 2: BlobPlot analysis of the 63 contigs in the chromosome-level assembly of *P. vulnus*, illustrating GC content distribution and taxonomic assignments based on BLAST hits to the NCBI nt database.** Three contigs were initially matched to Arthropoda, Bacteroidota, and Echinodermata, respectively. However, detailed examination of the aligned regions revealed that each contig contained only short stretches of a few hundred base pairs matching these taxa. Given the limited genomic data currently available for *P. vulnus* and the *Pratylenchus* genus as a whole in NCBI nt database, we consider it more appropriate to classify these sequences as no-hit.

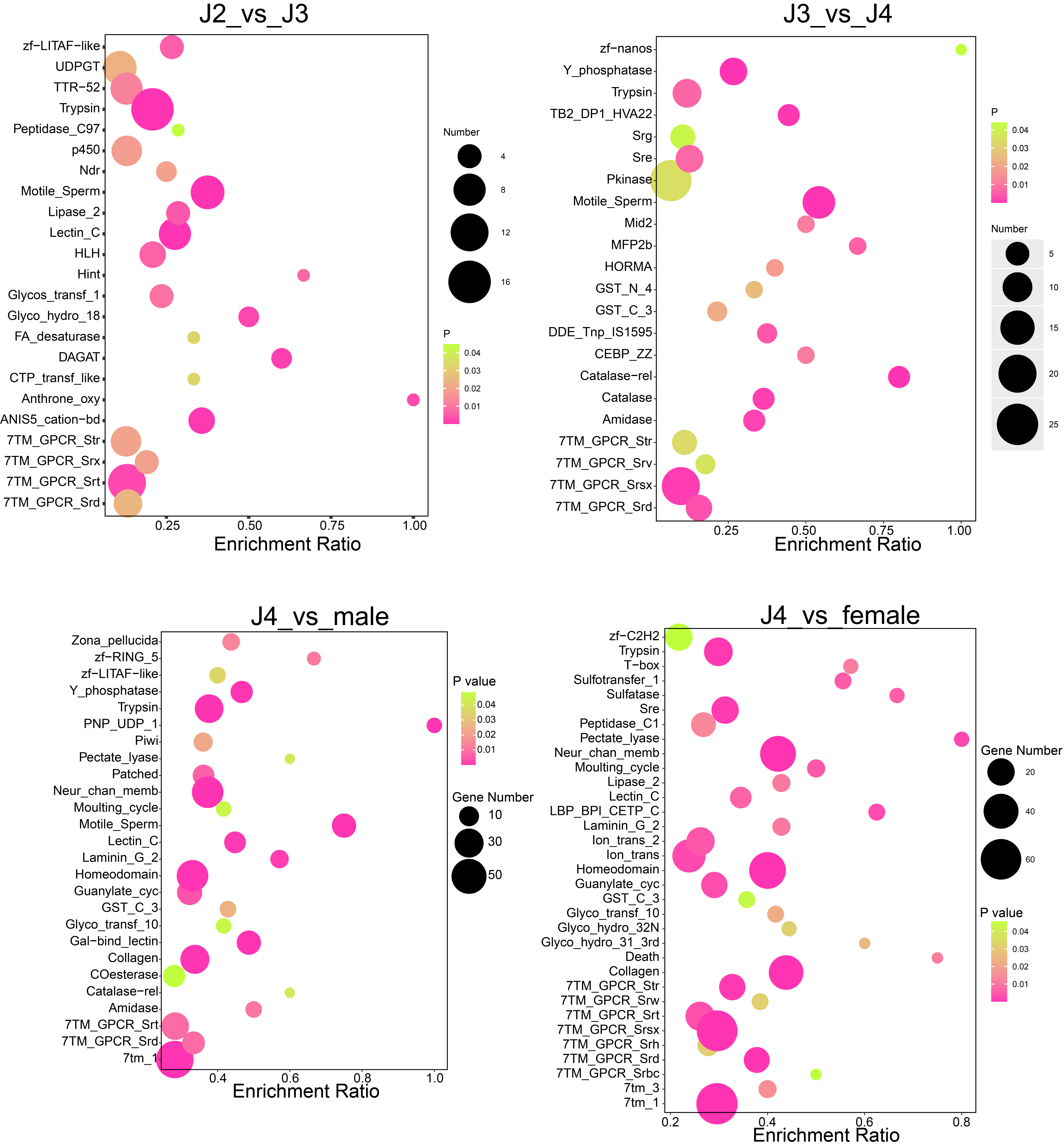

**Supplementary Fig. 3: Pfam functional enrichment analysis of differentially expressed genes (DEGs) across developmental stages and conditions in *P. vulnus*.** Bubble plots represent significantly enriched protein family (Pfam) domains among DEGs between the indicated comparisons: second-stage juveniles (J2) vs. J3, J3 vs. J4, J4 vs. males, J4 vs. females. Bubble size corresponds to the number of genes associated with each Pfam domain, while bubble color reflects statistical significance (p-value), with warmer colors indicating higher significance. Enrichment ratio on the x-axis indicates the proportion of observed to expected gene counts in each Pfam category.

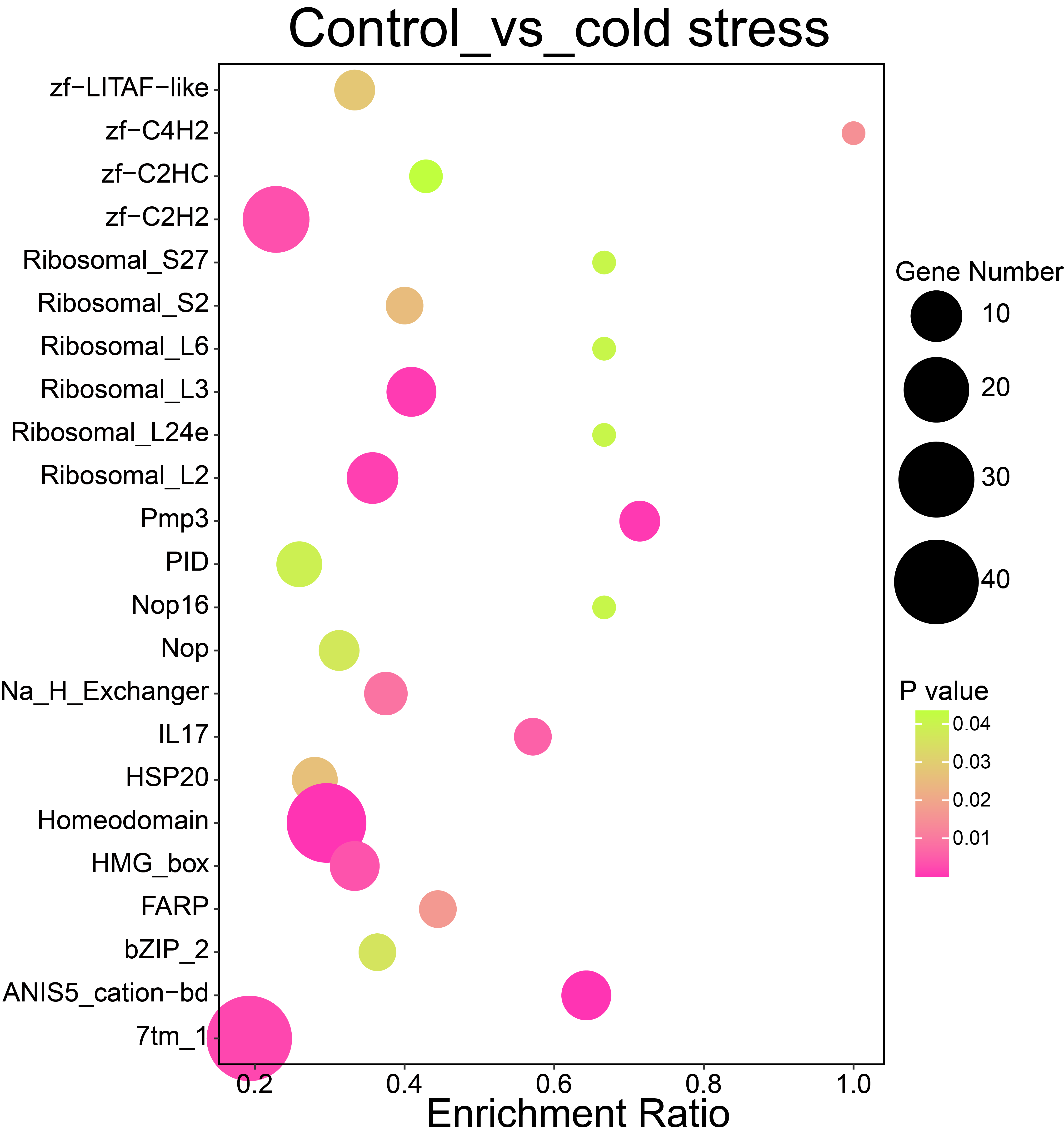

**Supplementary Fig. 4: Pfam functional enrichment analysis of DEGs between control and cold stress.** The control is the transcriptome data of the mix stage of the parasitic grapevine, and the cold-stress is also the mix stage of the parasitic grapevine, but it was treated with cold stress at 4°C for 6 hours.

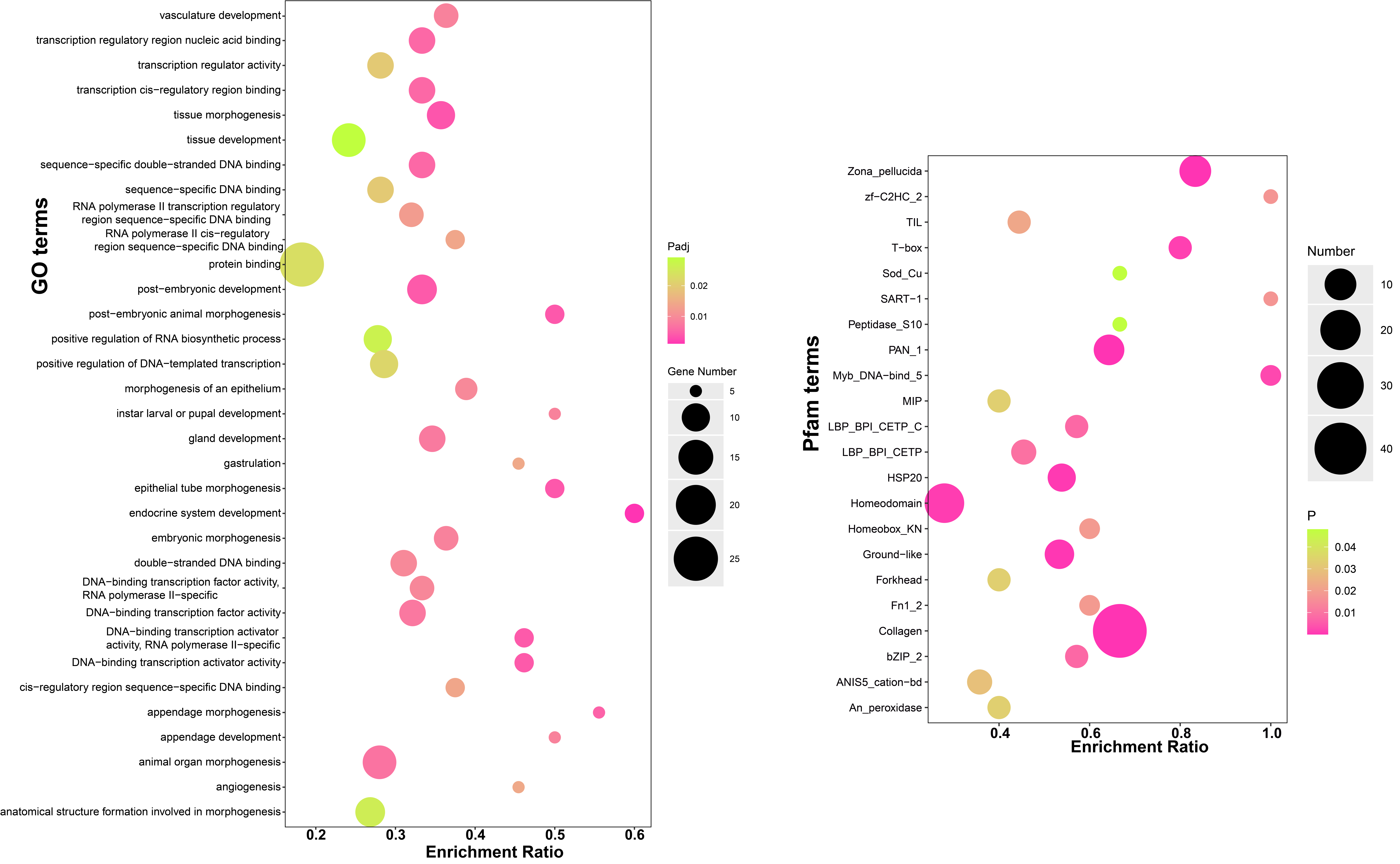

**Supplementary Fig. 5: Function enrichment analysis of DEGs upregulated in *P. vulnus* parasitizing walnut compared to the grapevine.**

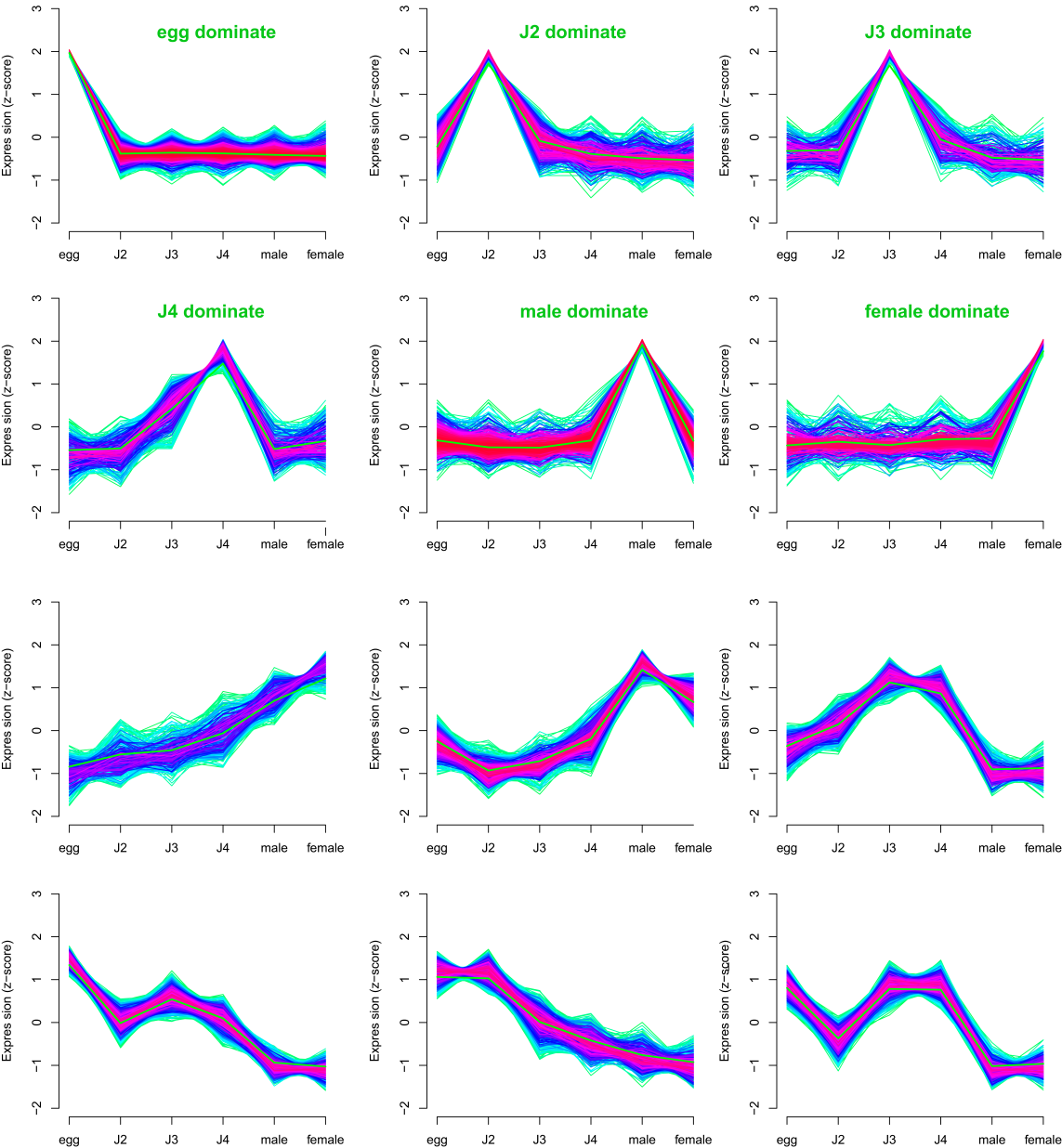

**Supplementary Fig. 6: Clustering analysis of gene expression profiles across developmental stages of *P. vulnus*.** Expression patterns are represented by Z-score transformed TPM (transcripts per million) values for each developmental stage (egg, J2, J3, J4, male, and female). The plots highlight distinct clusters of genes with dominant expression in specific developmental stages, indicating stage-specific gene regulation and functional specialization during nematode development.

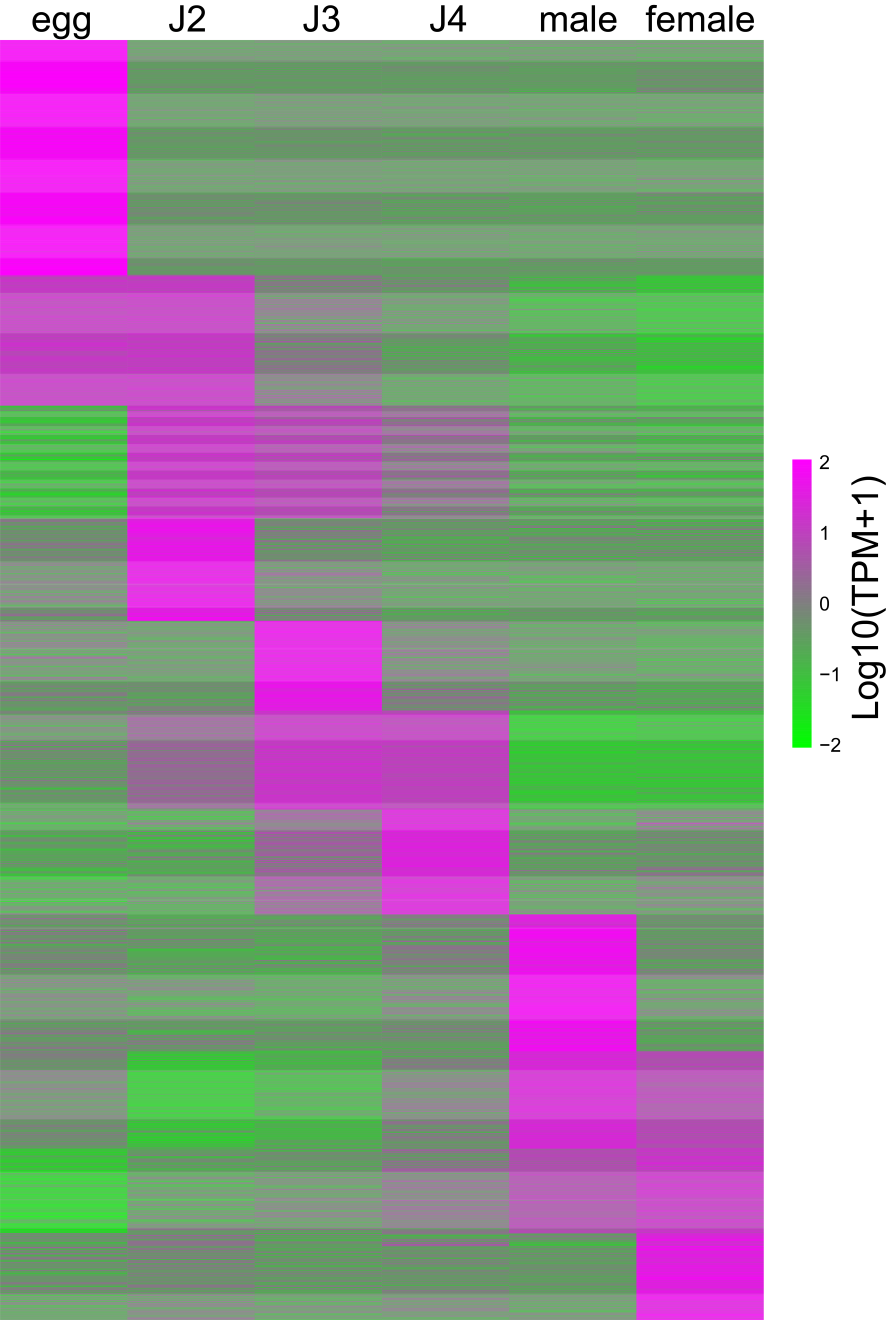

**Supplementary Fig. 7: Heatmap showing stage-specific dominant gene expression clusters across developmental stages of *P. vulnus*.** Gene expression values are represented as Log10(TPM + 1), with distinct clusters highlighting genes predominantly expressed at each stage (egg, J2, J3, J4, male, and female). High expressions are indicated in magenta, while low expressions are shown in green.

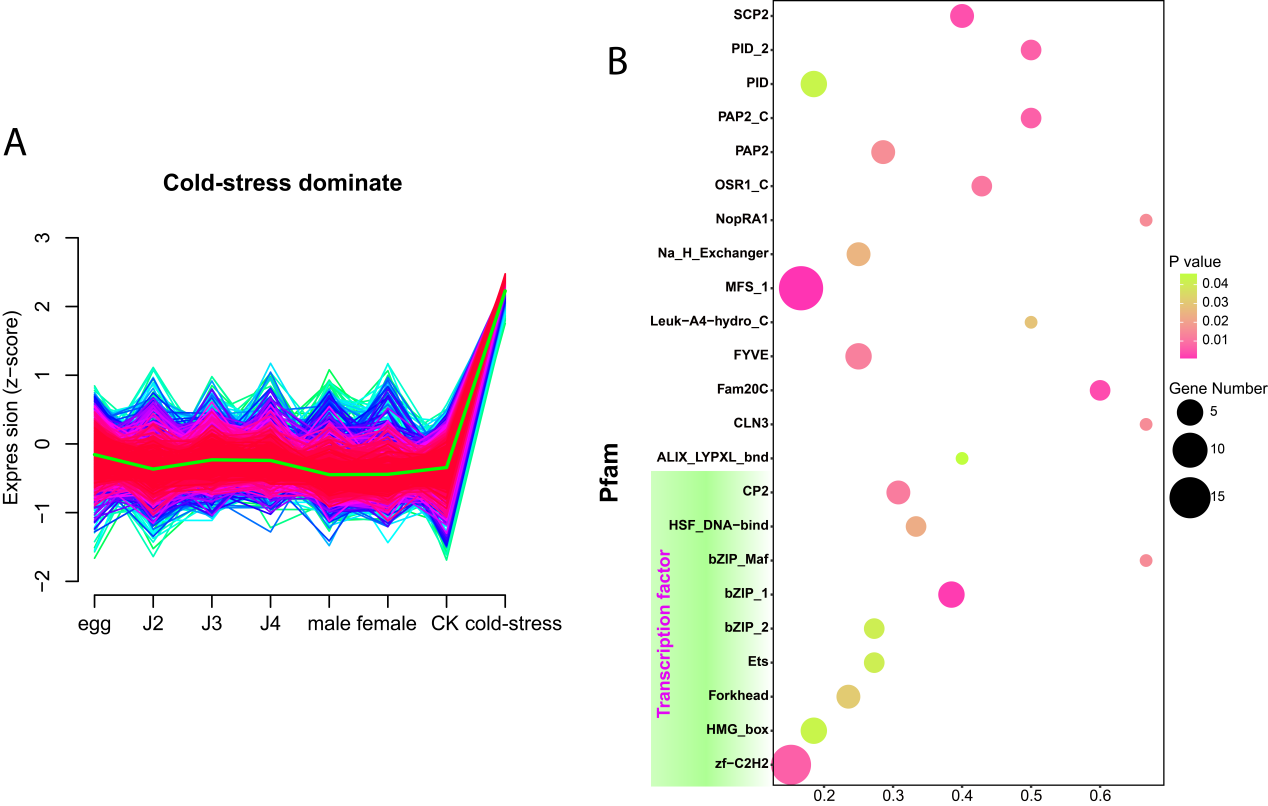

**Supplementary Fig. 8: Functional analysis of genes dominantly expressed under cold-stress conditions in *P. vulnus*.** (A) Mfuzz soft clustering analysis identified a distinct cluster of genes showing significantly increased expression under cold stress compared to other developmental stages and conditions. Gene expression values are normalized as Z-score. (B) Pfam domain enrichment analysis of genes dominantly expressed under cold stress. The bubble plot highlights significantly enriched Pfam domains, with bubble size indicating the number of genes within each domain and bubble color reflecting statistical significance (p-value).

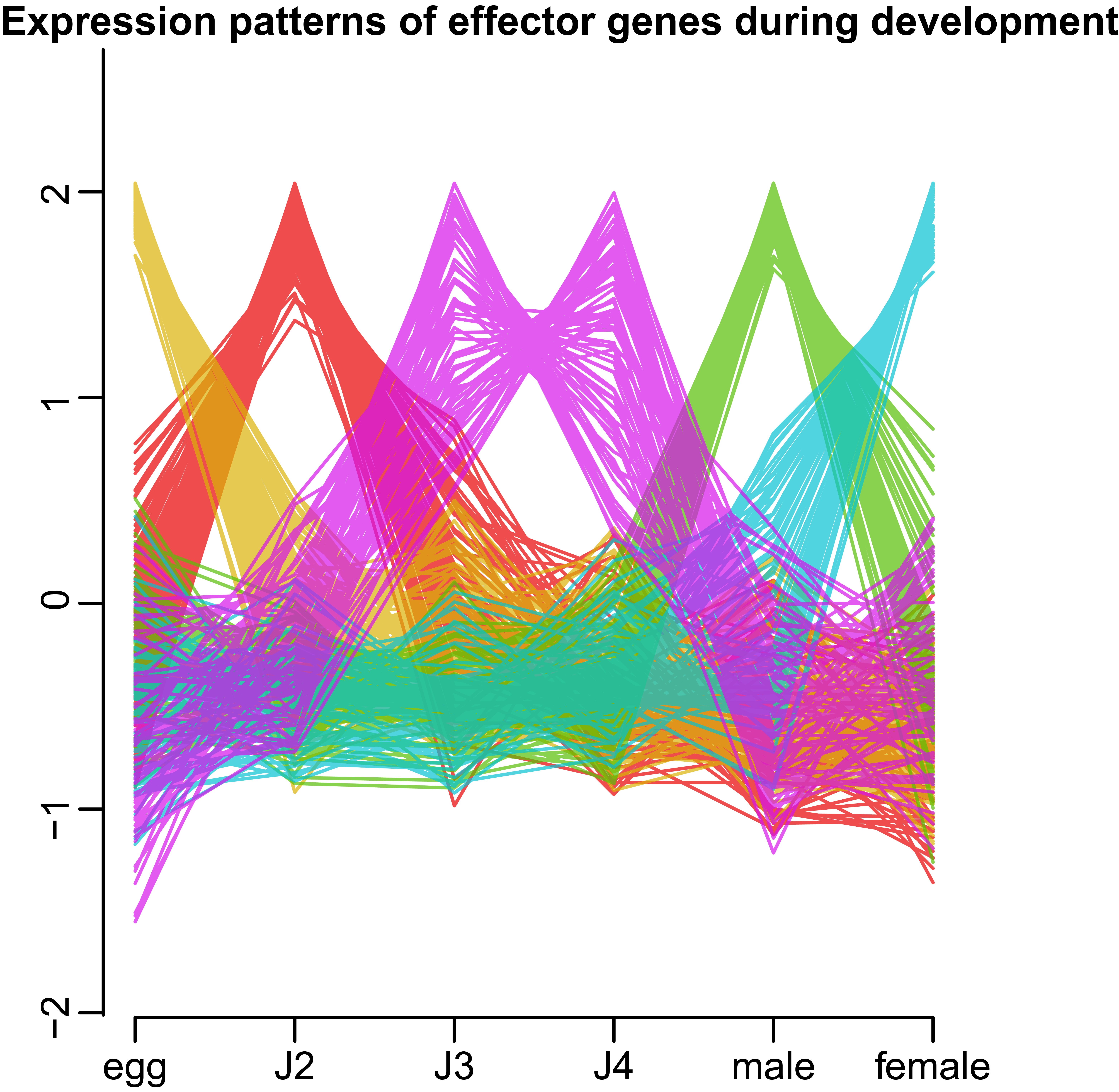

**Supplementary Fig. 9: Expression patterns of candidate effector genes at different stages of development.**

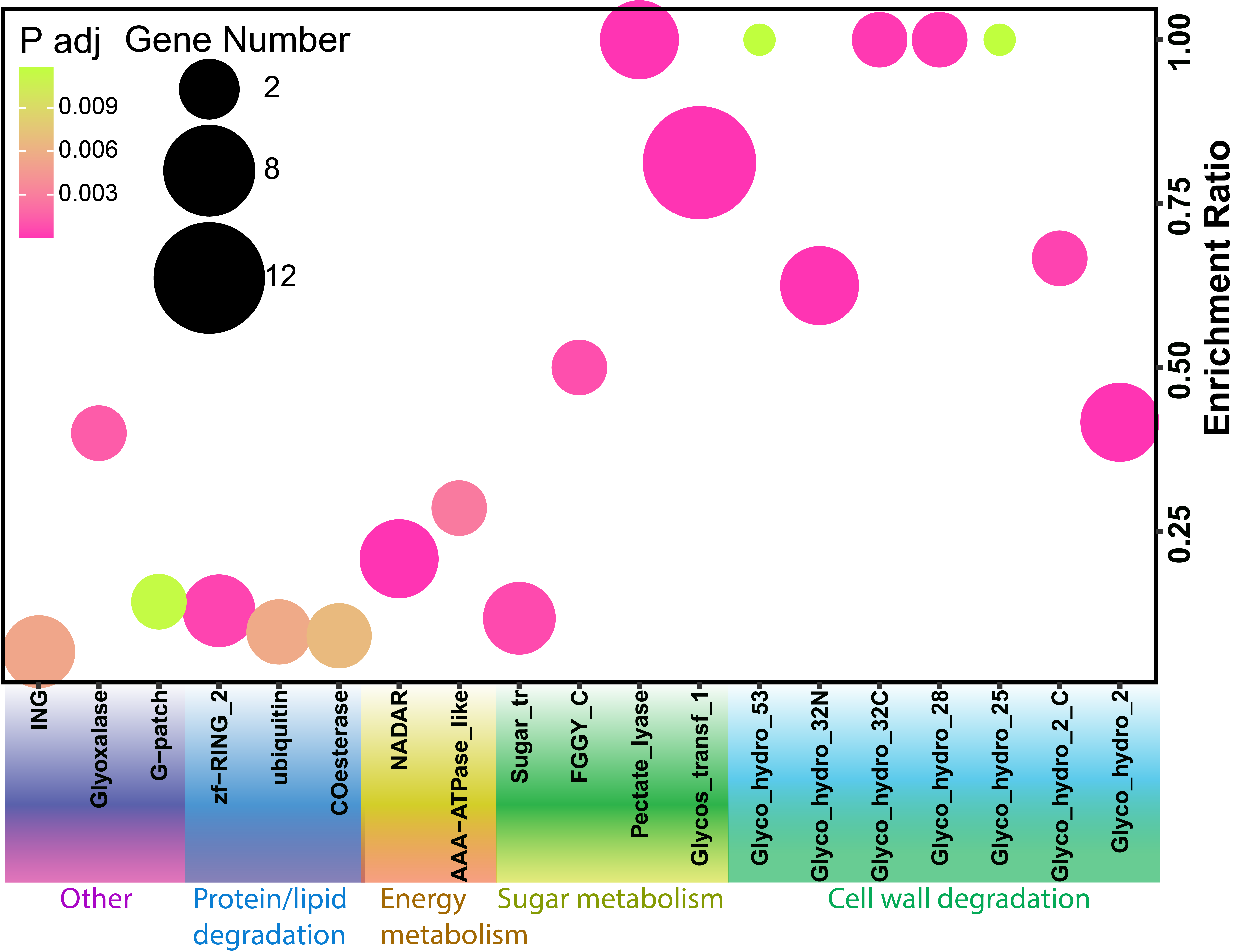

**Supplementary Fig. 10: Pfam domain enrichment analysis of proteins encoded by HGT-derived genes, highlighting protein families significantly enriched due to horizontal gene transfer events.**

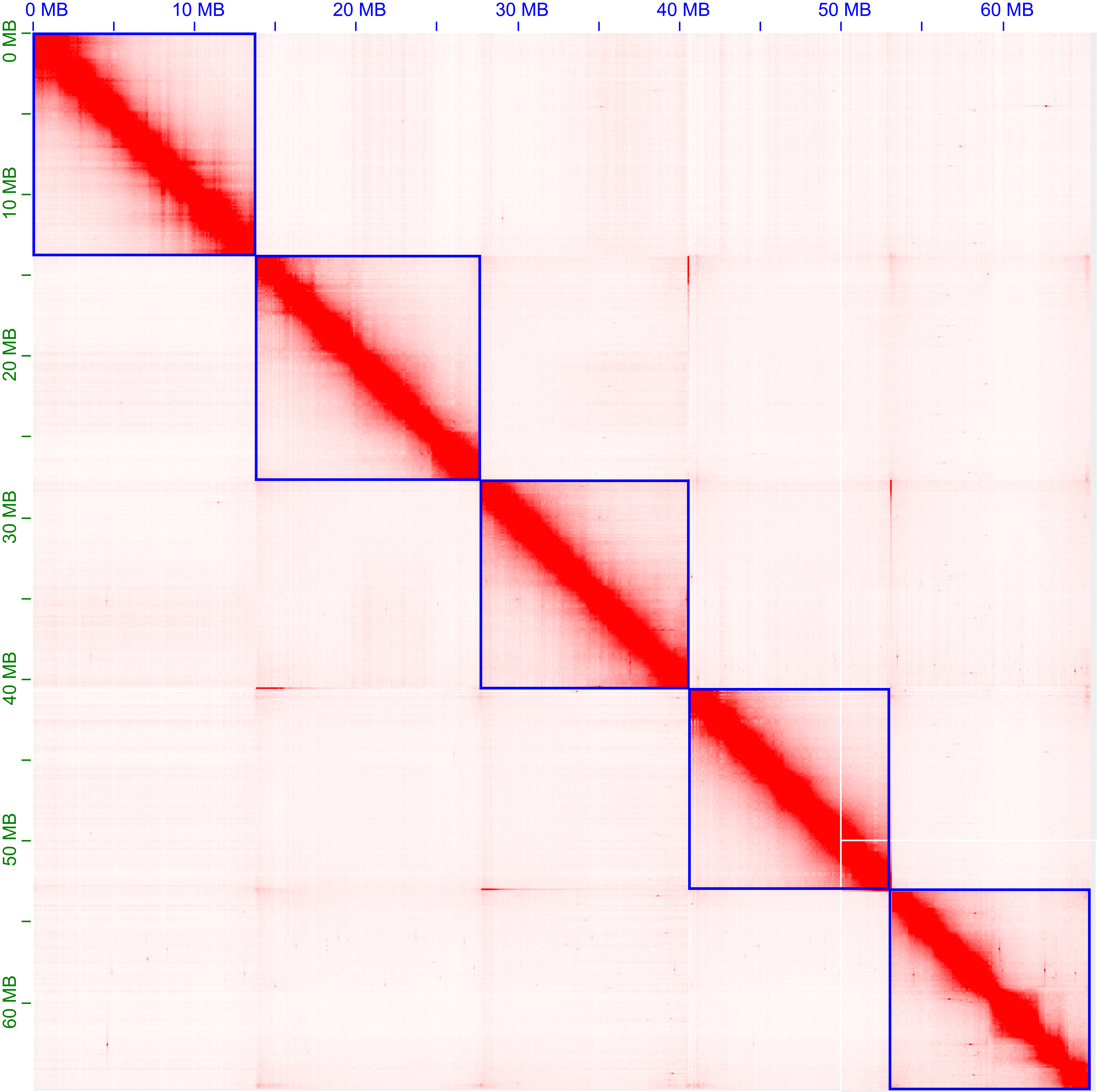

**Supplementary Fig. 11: After mounting the *Panagrellus* sp. LJ2284 contigs with Hi-C data, 5 chromosomes were obtained with a genome size of 65.4 Mb.** We downloaded the contig assembly of *Panagrellus* sp. LJ2284 and its Hi-C data, and then used these data to construct the genome at the chromosome level.

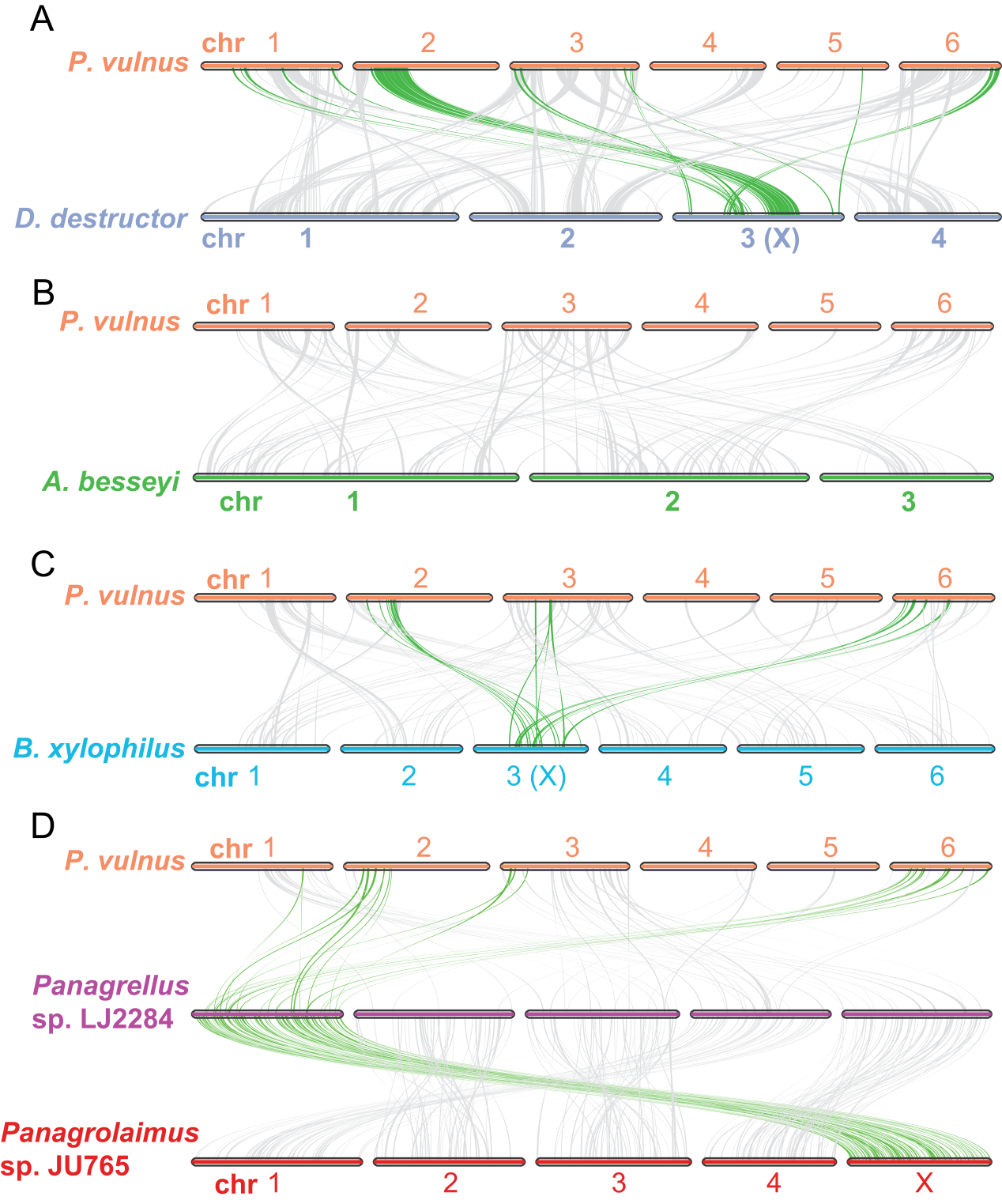

**Supplementary Fig. 12: Colinearity analysis of *P. vulnus* chromosomes compared to other nematode species based on protein sequence alignments.** (A) Colinearity between *P. vulnus* and *D. destructor*. The X chromosome of *D. destructor* is inferred based on Nigon element homology and established colinearity with *B. xylophilus*. (B) Colinearity between *P. vulnus* and *A. besseyi*. (C) Colinearity between *P. vulnus* and *B. xylophilus*. (D) Colinearity comparison among *P. vulnus*, *Panagrellus* sp. LJ2284, and *Panagrolaimus* sp. JU765. Chromosomes of *Panagrellus* sp. LJ2284 and *Panagrolaimus* sp. JU765 are highly conserved, showing no large-scale chromosomal rearrangements, whereas colinearity between these two species and *P. vulnus* is significantly lower, indicating extensive structural divergence.

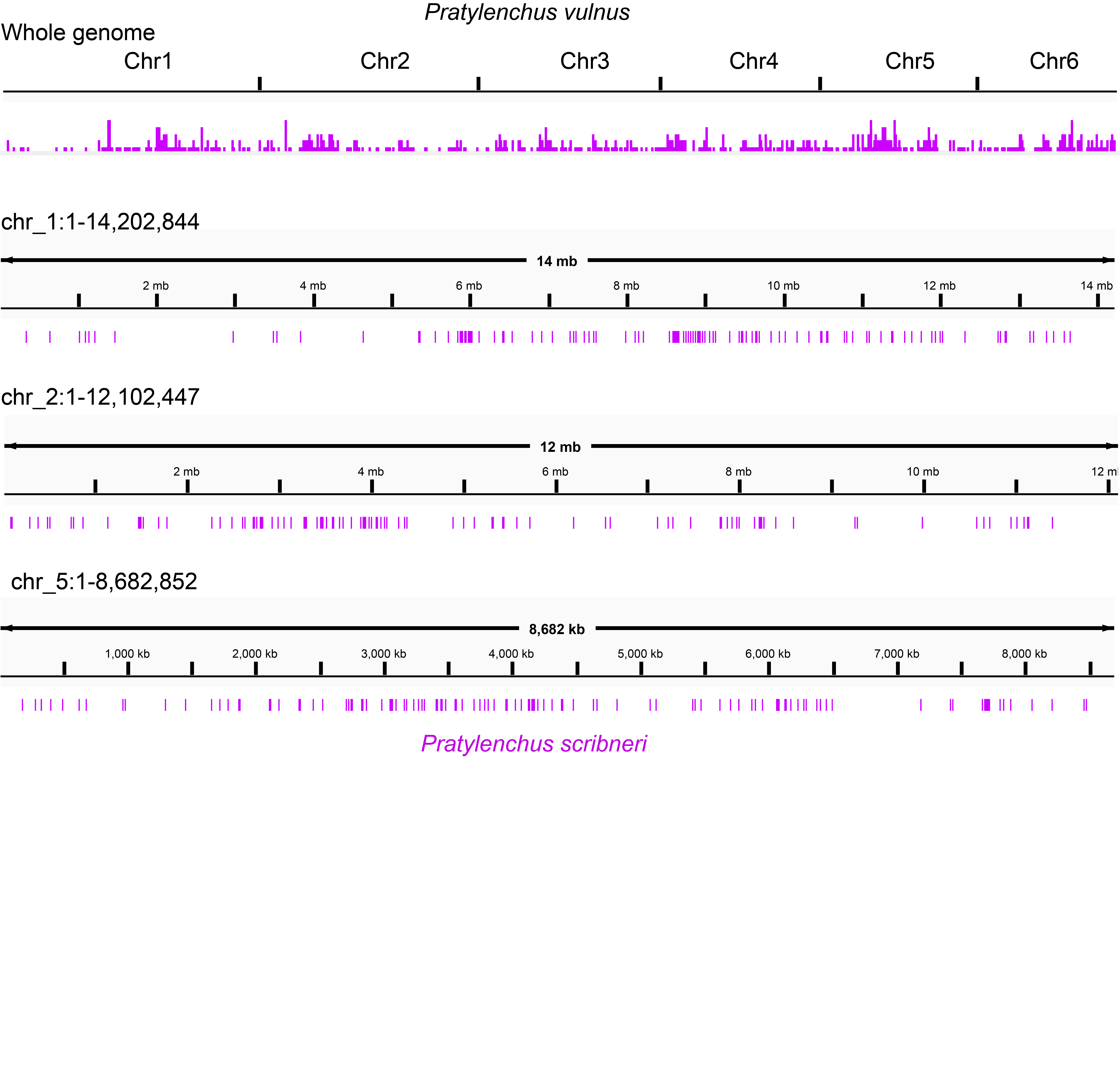

**Supplementary Fig. 13: Whole-genome sequence similarity comparison between *P. vulnus* and *P. scribneri*.** The draft genome of *P. scribneri* was aligned to the chromosome-level genome assembly of *P. vulnus*, and the aligned regions were converted into BED format. These homologous regions were visualized using IGV. In each track, the black at the top represent the *P. vulnus* genome, and the purple bars below indicate the aligned *P. scribneri* genome sequences.

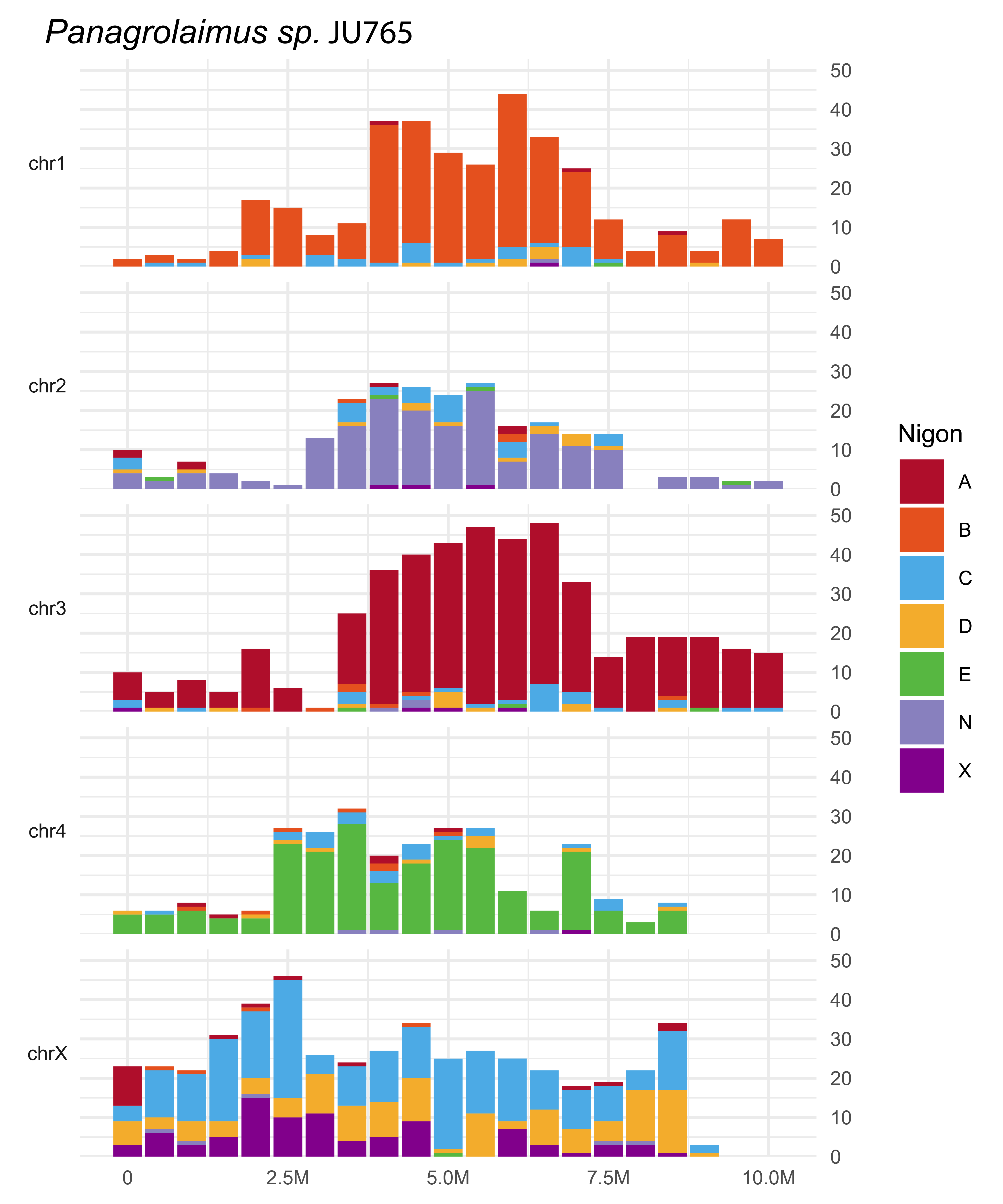

**Supplementary Fig. 14: The distribution of Nigon elements on the chromosome of *Panagrolaimus sp.* JU765.**

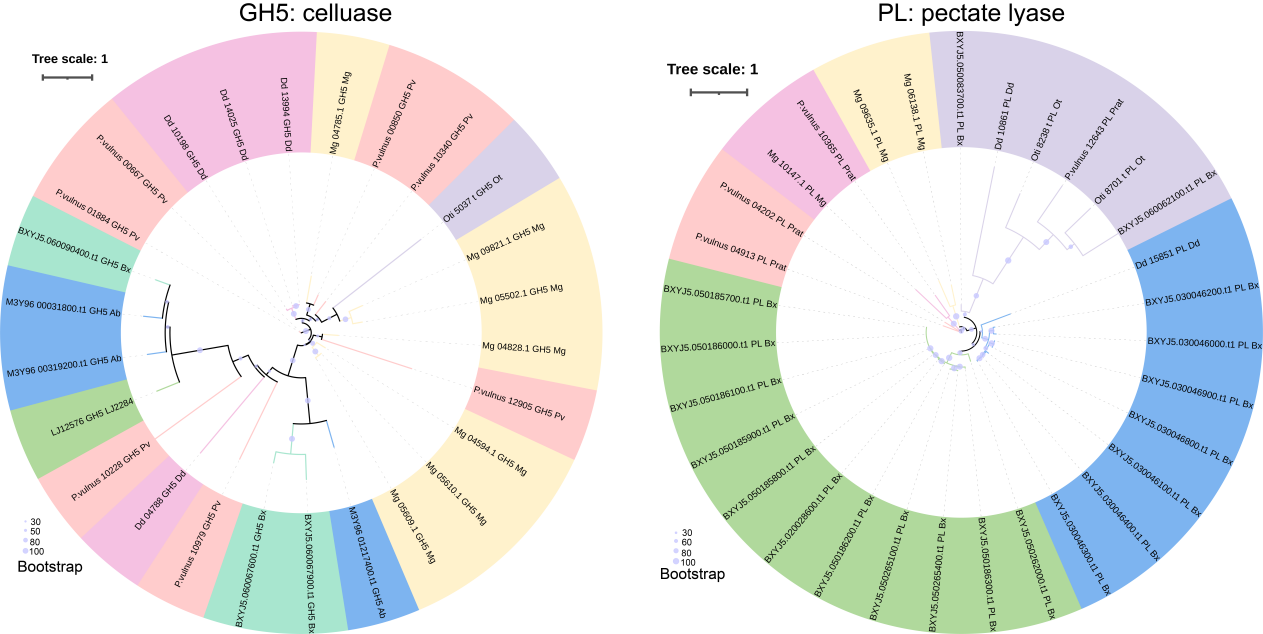

**Supplementary Fig. 15: Phylogenetic analysis of GH5 (cellulase) and PL (pectate lyase) gene families from eight nematode species.** Phylogenetic trees were constructed using protein sequences from GH5 cellulases (left) and PL pectate lyases (right). Branch colors correspond to different clades, and bootstrap values are indicated by branch thickness and opacity, with thicker branches representing higher support. These analyses highlight evolutionary relationships and potential gene duplication or diversification events within GH5 and PL families across nematodes.

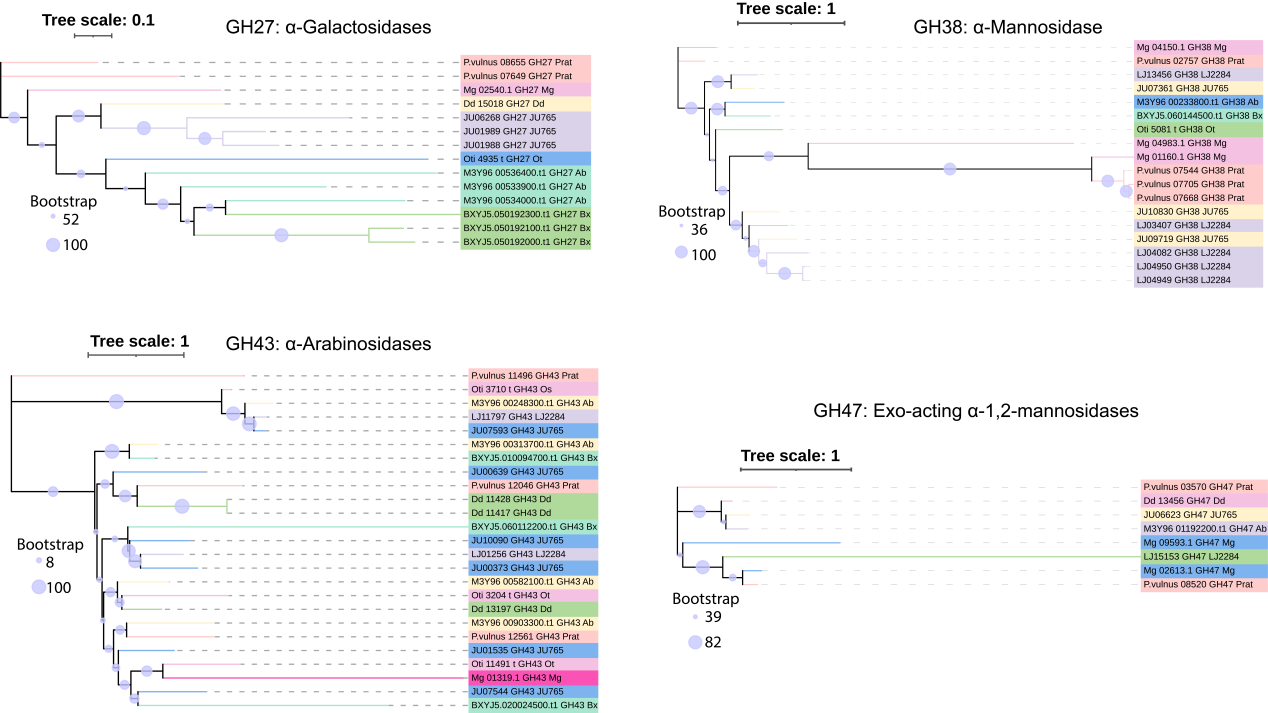

**Supplementary Fig. 16: Phylogenetic relationships of GH27 (α-galactosidases), GH38 (α-mannosidases), GH43 (α-arabinosidases), and GH47 (exo-acting α-1,2-mannosidases) gene families among eight nematode species analyzed in this study.** Trees illustrate evolutionary relationships based on protein sequences. Bootstrap support is indicated by circle size at the nodes, with larger circles representing higher confidence. Different branch colors represent distinct clades. The analysis highlights evolutionary divergence, conservation, and potential functional specializations of these glycoside hydrolase families across nematodes.

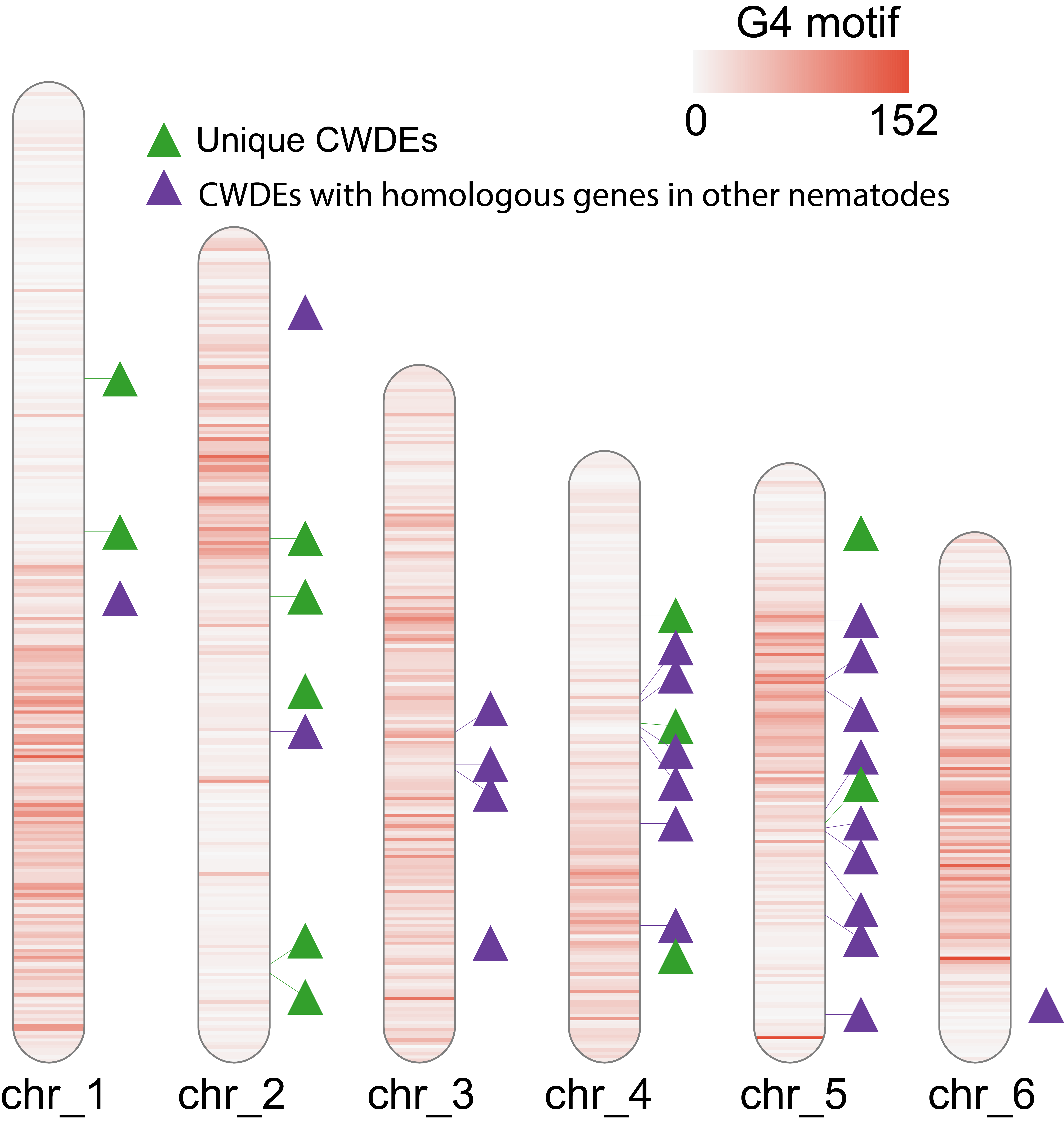

**Supplementary Fig. 17: Many of the phylogenetically distinct CWDE genes in *P. vulnus* are located in non-syntenic regions on chromosomes 1, 2, 4, and 5.**

**Supplementary Tables:**

**Supplementary Table 1 Summary of the TE in *P. vulnus* genome**

| Class | Count | bpMasked | %masked |
| --- | --- | --- | --- |
| DNA | | | |
| DTA | 6710 | 929916 | 1.51% |
| DTC | 852 | 143160 | 0.23% |
| DTH | 194 | 46745 | 0.08% |
| DTM | 9921 | 1706071 | 2.77% |
| Helitron | 2203 | 389578 | 0.63% |
| MULE-MuDR | 13 | 1582 | 0.00% |
| Maverick | 44 | 44320 | 0.07% |
| Merlin | 105 | 27160 | 0.04% |
| TcMar-Mariner | 99 | 62804 | 0.10% |
| TcMar-Tc1 | 30 | 25873 | 0.04% |
| LINE | | | |
| CR1 | 273 | 42765 | 0.07% |
| LTR | | | |
| Copia | 27 | 1559 | 0.00% |
| ERV1 | 109 | 7597 | 0.01% |
| Gypsy | 67 | 7429 | 0.01% |
| unknown | 2 | 103 | 0.00% |
| MITE | | | |
| DTA | 5332 | 549216 | 0.89% |
| DTC | 184 | 22481 | 0.04% |
| DTH | 180 | 19963 | 0.03% |
| DTM | 2015 | 183118 | 0.30% |
| Other | | | |
| Helitron | 7 | 872 | 0.00% |
| SINE 5S | 31 | 4851 | 0.01% |
| Unknown | 10252 | 1621536 | 2.63% |
| total interspersed | 38650 | 5838699 | 9.46% |
| Simple_repeat | 428 | 78830 | 0.13% |
| **Total** | **39078** | **5917529** | **9.59%** |

**Supplementary Table 2 Summary of CWDEs of different nematodes**

| Family | Ot | JU765 | LJ2284 | Bx | Ab | Dd | Pv | Mg |
| --- | --- | --- | --- | --- | --- | --- | --- | --- |
| GH3 | 1 | 0 | 4 | 2 | 0 | 1 | 1 | 0 |
| GH5 | 1 | 0 | 1 | 3 | 3 | 4 | 7 | 7 |
| GH45 | 1 | 0 | 0 | 9 | 4 | 0 | 0 | 0 |
| GH27 | 1 | 3 | 0 | 3 | 3 | 1 | 2 | 1 |
| GH31 | 16 | 28 | 8 | 5 | 8 | 7 | 3 | 0 |
| GH35 | 0 | 3 | 2 | 0 | 2 | 1 | 1 | 0 |
| GH38 | 1 | 3 | 5 | 1 | 1 | 0 | 4 | 3 |
| GH43 | 3 | 6 | 2 | 3 | 4 | 3 | 3 | 1 |
| GH47 | 0 | 1 | 1 | 0 | 1 | 1 | 2 | 2 |
| CBM13 | 9 | 15 | 18 | 48 | 7 | 7 | 13 | 3 |
| GH18 | 10 | 8 | 10 | 7 | 4 | 5 | 3 | 0 |
| GH19 | 0 | 1 | 1 | 1 | 1 | 0 | 1 | 0 |
| GH20 | 3 | 1 | 3 | 2 | 2 | 0 | 1 | 0 |
| PL | 2 | 0 | 0 | 20 | 0 | 2 | 5 | 3 |
| GH16 | 8 | 8 | 7 | 26 | 11 | 10 | 10 | 1 |
| GH2 | 2 | 2 | 5 | 1 | 2 | 2 | 2 | 2 |
| GH15 | 1 | 0 | 0 | 0 | 0 | 0 | 0 | 0 |
| GH25 | 6 | 9 | 5 | 11 | 2 | 0 | 3 | 4 |
| GH32 | 6 | 11 | 5 | 8 | 7 | 4 | 5 | 1 |
| GH56 | 0 | 2 | 1 | 2 | 1 | 0 | 0 | 0 |
| Total | 71 | 101 | 78 | 152 | 63 | 48 | 66 | 28 |

Ot: *O. tipulae*; JU765: *Panagrolaimus* sp. JU765; LJ2284: *Panagrellus* sp. LJ2284; Bx: *B. xylophilus*; Ab: *A. besseyi*; Dd: *D. destructor*; Pv: *P. vulnus*; Mg: *M. graminicola*
